# Convergent mutations in tissue-specific regulatory regions reveal novel cancer drivers

**DOI:** 10.1101/2020.08.21.239954

**Authors:** Nasa Sinnott-Armstrong, Jose A. Seoane, Richard Sallari, Jonathan K. Pritchard, Christina Curtis, Michael P. Snyder

## Abstract

Although much effort has been devoted to identifying coding mutations across cancer types, regulatory mutations remain poorly characterized. Here, we describe a framework to identify non-coding drivers by aggregating mutations in cell-type specific regulatory regions for each gene. Application of this approach to 2,634 patients across 11 human cancer types identified 60 pan-cancer, 22 pan-breast and 192 cancer specific candidate driver genes that were enriched for expression changes. Analysis of high-throughput CRISPR knockout screens revealed large, cancer specific growth effects for these genes, on par with coding mutations and exceeding that for promoter mutations. Amongst the five candidate drivers selected for further analysis, four (*IPO9, MED8, PLEKHA6*, and *OXNAD1)* were associated with survival across multiple cancer types. These studies demonstrate the power of our cell-type aware, convergent regulatory framework to define novel tissue specific cancer driver genes, considerably expanding evidence of functional non-coding mutations in cancer.

## Introduction

To date, much effort has been devoted to the analysis of coding regions within the human genome to define somatic alterations associated with tumor growth and progression (Bailey et al., 2018; Lawrence et al., 2014; Zehir et al., 2017). While many recurrent clonal coding mutations have been defined, non-coding elements (including promoters and enhancers) implicated in malignancy have been far more elusive due to the need for large cohorts with whole genome sequencing (WGS) data and new analytic approaches. Indeed, attempts to locate regulatory elements enriched for functional mutations (Araya et al., 2016; Feigin et al., 2017; Melton et al., 2015; Weinhold et al., 2014; Zhu et al., 2020) have revealed only a handful of target genes, most of which are associated with core promoter variants. An example is the canonical oncogene *TERT*, where promoter mutations can induce c-Myc activation and telomeric immortalization (Berger et al., 2012; Huang et al., 2013; Wu et al., 1999). However, the vast majority of genes are regulated by promoters as well as proximal and distal enhancer elements (Schmidt et al., 2010), suggesting that the latter may harbor as of yet undiscovered mutations. Indeed, the long non-coding RNA (lncRNA) gene *PVT1* was recently identified as a tissue-specific tumor suppressor DNA boundary element that regulates *MYC* transcription (Cho et al., 2018), demonstrating a role for regulatory sequences of lncRNAs in malignancy. A recent paper by Rheinbay et al identified a small number (4–5) of driver mutations when combining coding and non-coding genomic elements per cancer genome. However, even in this most recent study, analyses suggest that discovery of noncoding mutations and driver genes is far from complete (Rheinbay et al., 2020).

The tissue-specific epigenomic landscape of a cell dictates its response to oncogenic cues and influences the selection of somatic alterations during tumor initiation (Lawrence et al., 2014; Lowdon and Wang, 2017; Sack et al., 2018). Accordingly, we reasoned that tissue-specific annotations may increase the power and interpretability of cancer driver gene discovery. As evidenced by their enrichment in genome-wide association studies (GWAS), expression quantitative trait loci (eQTLs), and cross-species conservation analyses, sequence alterations in regulatory elements are associated with functional changes in the expression of downstream target genes and disease phenotypes (Maurano et al., 2012; Schaub et al., 2012; Zhou et al., 2020). Meanwhile, putative regulatory element mutations have been shown to affect cancer driver gene expression in relevant tissues (Takeda et al., 2018; Zhang et al., 2018). Therefore, the systematic analysis of regulatory variants within active elements of the corresponding cell type of origin may improve the power to detect non-coding cancer associated mutations.

Here, we leverage these principles to develop a generalizable analytic framework to characterize cell-type-specific regulatory landscapes and non-coding mutational burden across 2,634 patients spanning 11 cancer types (Supplemental Table 1). We focused on regulatory variants within active elements in the cell type of origin, defined by the chromatin state of the corresponding *enhancer* or *promoter*. To increase the power to detect disease-associated variants, we aggregated regulatory information across all elements for each gene, similar to recent work examining the *ESR1* locus in breast cancer (Bailey et al., 2016) and prostate cancer (Sallari et al., 2017). Using this approach, we found both known and novel recurrently mutated regulatory regions, the majority of which were associated with dysregulated expression of nearby genes and differential survival outcomes. In particular, we identify *IPO9* as a novel regulatory driver mutation in breast cancer. Using high-throughput CRISPR screen data across cancer cell lines (Meyers et al., 2017), we demonstrate that genes harboring recurrent regulatory mutations, including *IPO9, GUK1, MED8*, and *OXNAD1*, were associated with larger *in vitro* growth effects on average than genes enriched for coding mutations. Together, these results highlight the power of aggregating regulatory information and the use of cell-type-aware models to define novel oncogenic drivers across diverse cancers.

## Results

### Analytical Framework

We reasoned that the power to discover novel regulatory regions as well as driver genes would be improved by combining regulatory information for each gene, analogous to burden tests aggregating exonic information for coding sequences (Figure 1A). In order to capture information relevant for each cancer type, we used cell type specific epigenetic data available from the ENCODE and Roadmap Epigenome projects. We estimated mutational enrichment within regulatory regions of each gene by permutation testing (Methods). To implement this approach, we first linked the distal enhancer elements defined by the Roadmap Epigenomics Consortium (Roadmap Epigenomics Consortium et al., 2015) to each of the 18,729 GENCODE genes using the correlation-based links from Roadmap (Figure 1B). Each distal element can be assigned to one or more genes. To verify the quality of these enhancer-promoter links, we counted the number of linked genes present at each enhancer element (Supplemental Figure 1C). Each distal element linked to ∼5 genes on average, consistent with other studies (Fishilevich et al., 2017).

**Figure 1:**
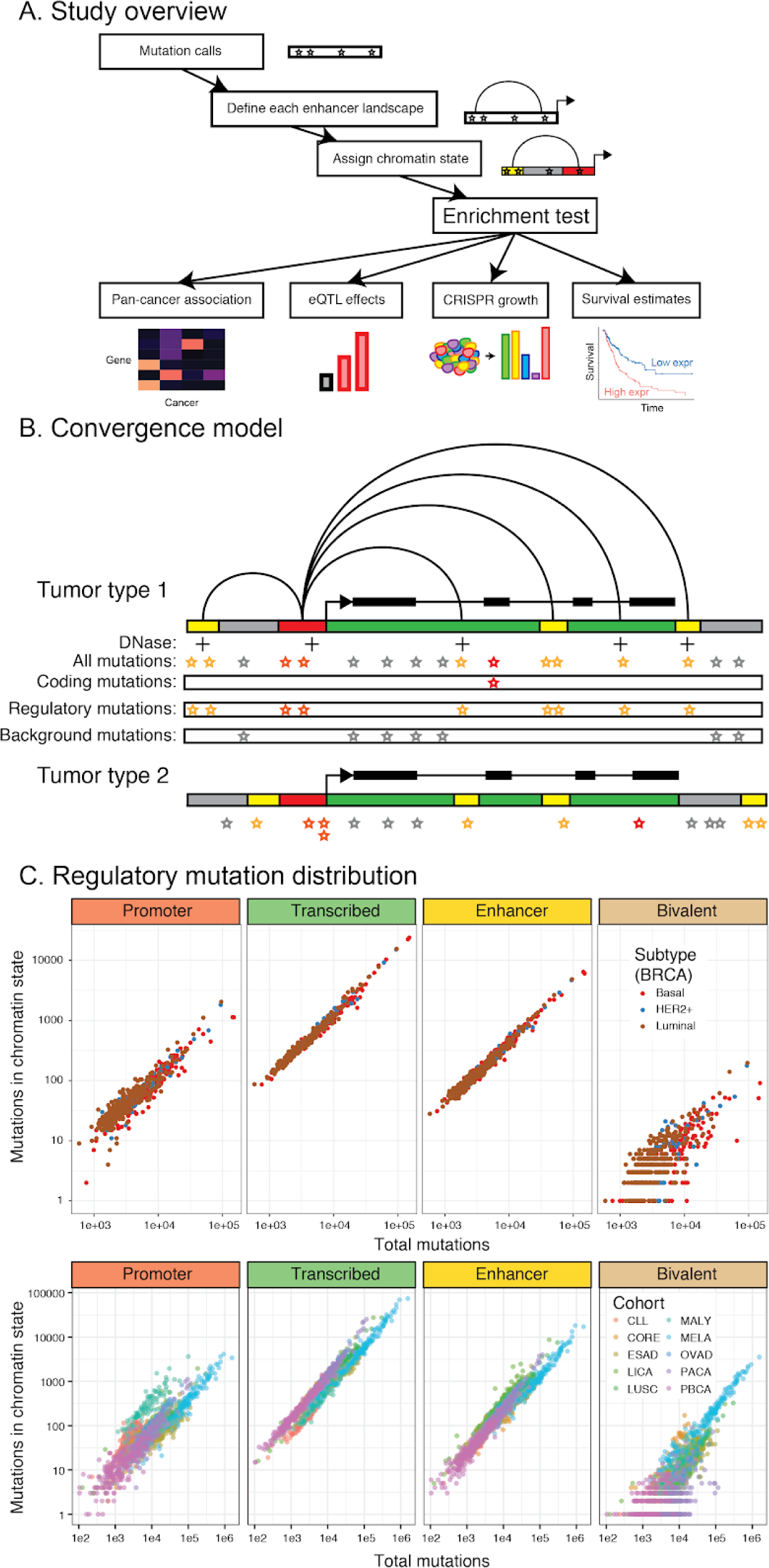
Model for aggregating mutations in gene-associated regulatory regions. **A**.**Study overview**. Overview of approach to evaluate aggregate mutational burden in non-coding regulatory regions across cancer types, their functional effects, and clinical outcome associations. **B**.**Convergence model**. Mutations accumulate in coding sequences and promoters, as detected in existing methods, but non-promoter regulatory mutations are likely spread across enhancer elements. Jointly testing specific regulatory regions can therefore increase the signal of mutational burden at a given gene, similar to an exome burden test. Both mutations and regulatory annotations change between tumor types. **C**.**Regulatory mutation distribution**. Ordered distribution of mutation counts per individual for each of the cancer types studied in active and bivalent chromatin state annotations. (x) axis total mutations for a given tumor, and (y) axis number of mutations in a given chromatin state (promoter, enhancer, transcribed, or bivalent) for this tumor. Each point represents a single tumor within each subplot. For breast cancer, the three subtypes analysed separately are individually plotted.

To assess the quality of our regulatory links, we next intersected these links with chromatin states from the corresponding cell type, producing a canonical enhancer-enriched distribution of regulatory activity (Supplemental Figure 1D, Supplemental Table 2). We compared the chromatin state annotations within each cancer type on each side of a regulatory link and discovered an enrichment of repressed regulatory elements linked to repressed promoters and active regulatory elements linked to active promoters, consistent with expectations of domain-level activation (Rao et al., 2014) (Supplemental Figure 1E). These results indicate that both chromatin states and enhancer-gene links are stable and high quality.

To evaluate mutational enrichment in regulatory regions of all genes, we used SNV and indel calls from WGS data from the International Cancer Genome Consortium (ICGC) focusing on 11 cancer types with a minimum of n=90 individuals and tissue matched epigenetic data (Figure 1C, Supplemental Table 1). In addition, the breast cancer cohort was sufficiently large to enable evaluation of the etiologically distinct Basal, Luminal, and HER2+ subgroups (Nik-Zainal et al., 2016). The variants from each cohort were normalized for regional patient mutation rate (Methods), chromatin state, and cancer type, intersected with each gene’s aggregated regulatory regions and evaluated for mutational enrichment. Enrichment was assessed by permutation testing (as in (Sallari et al., 2017); 5000+ iterations), where a matching background set of regulatory elements were randomly assigned to each gene (maintaining mutation rate and chromatin state) and the number of mutations scored (Methods).

### Excess mutational burden in aggregate distal regulatory regions in breast cancer

We first evaluated this approach in a WGS dataset composed of 560 breast cancers stratified by three major subtypes: Basal (n = 167), Luminal (n = 320), and HER2+ (n = 73) (Nik-Zainal et al., 2016). We performed enrichment tests on 57,534 FANTOM-derived promoters for 20,209 Ensembl-annotated genes, where promoters for the same gene were concatenated when evaluating enrichments (Methods). Consistent with previous results, we observed an enrichment in mutations in the shared promoter of *RMRP* and *CCDC107* across the individual breast cancer cohorts (Nik-Zainal et al., 2016; Rheinbay et al., 2017). Combining p-values across the three breast cancer subtypes via Fisher’s method revealed enrichment of promoter mutation in *TP53* and *CCDC107*, as previously reported. When considering only active promoter elements, we identify enrichments in *WDR74, ZNF143, MFSD11, SRSF2, VMA21, CDC42BPB*, and *TMEM189* (Supplemental Table 3). Thus, analysis of single regulatory elements reveals excess mutational burden in numerous previously identified drivers, as well as novel candidate drivers.

We hypothesized that aggregating distal regulatory elements would yield increased power to detect candidate driver genes. For each of the 18,729 GENCODE genes we aggregated the promoter-interacting regulatory elements and tested for an excess or overburdening of distal mutations. In order to resolve cell-type-specific effects, we examined combinations of different chromatin states that represent the regulatory profile of mammary epithelial cells (e.g. poised enhancers, active enhancers, promoters, Supplemental Table 2). Using this approach, we identify 22 putative distal regulatory driver genes with FDR < 10%, spanning numerous regulatory states. These candidates included known driver genes such as *MSL3* (Leiserson et al., 2013) *and HLE* (Osborne et al., 2010) (Supplemental Table 4). In addition, we found significant enrichment for mutations in regulatory regions of 17 novel genes, most notably *IPO9*, which was specifically enriched in enhancer marked chromatin (Figure 2C). Mutations in regulatory regions of *IPO9* were significantly overburdened in basal subtype tumors where 15 patients harbored 16 mutations, compared to an expectation of ∼3.6 patients (4.2-fold enrichment, permutation p-value < 3.2e-6, Methods). An additional 3 patients across the other subgroups exhibited IPO9 mutations, bringing the total to 18 (Fisher combined, FDR adjusted q-value across all three breast cancer subtypes = 0.068). Additionally, *PYCR2* exhibited an excess of regulatory mutations (23 mutations across 22 patients, q-value = 0.002) in active promoter & strong enhancer (H3K4me3)-marked regions, as did *SDE2* (18 mutations across 17 patients, q-value = 0.023), *SRP9* (24 mutations in 23 patients, q-value = 0.02), and *PLEKHA6* (22 mutations in 21 patients, q-value = 0.04, Supplemental Figure 2C). *PYCR2* catalyzes the last step of proline synthesis from glutamate in the mitochondrion (De Ingeniis et al., 2012); *SDE2* is a telomere repair gene implicated in cell cycle regulation (Jo et al., 2016); *SRP9* binds and inhibits *Alu* element translation (Chang et al., 1996); and *PLEKHA6* is poorly characterized. Also of note, luminal tumors comprise a heterogeneous group that can be stratified based on genomic features (Rueda et al., 2019), hence it is not surprising that mutational enrichment is weaker than observed in Basal and HER2+ tumors (Figure 2D).

**Figure 2:**
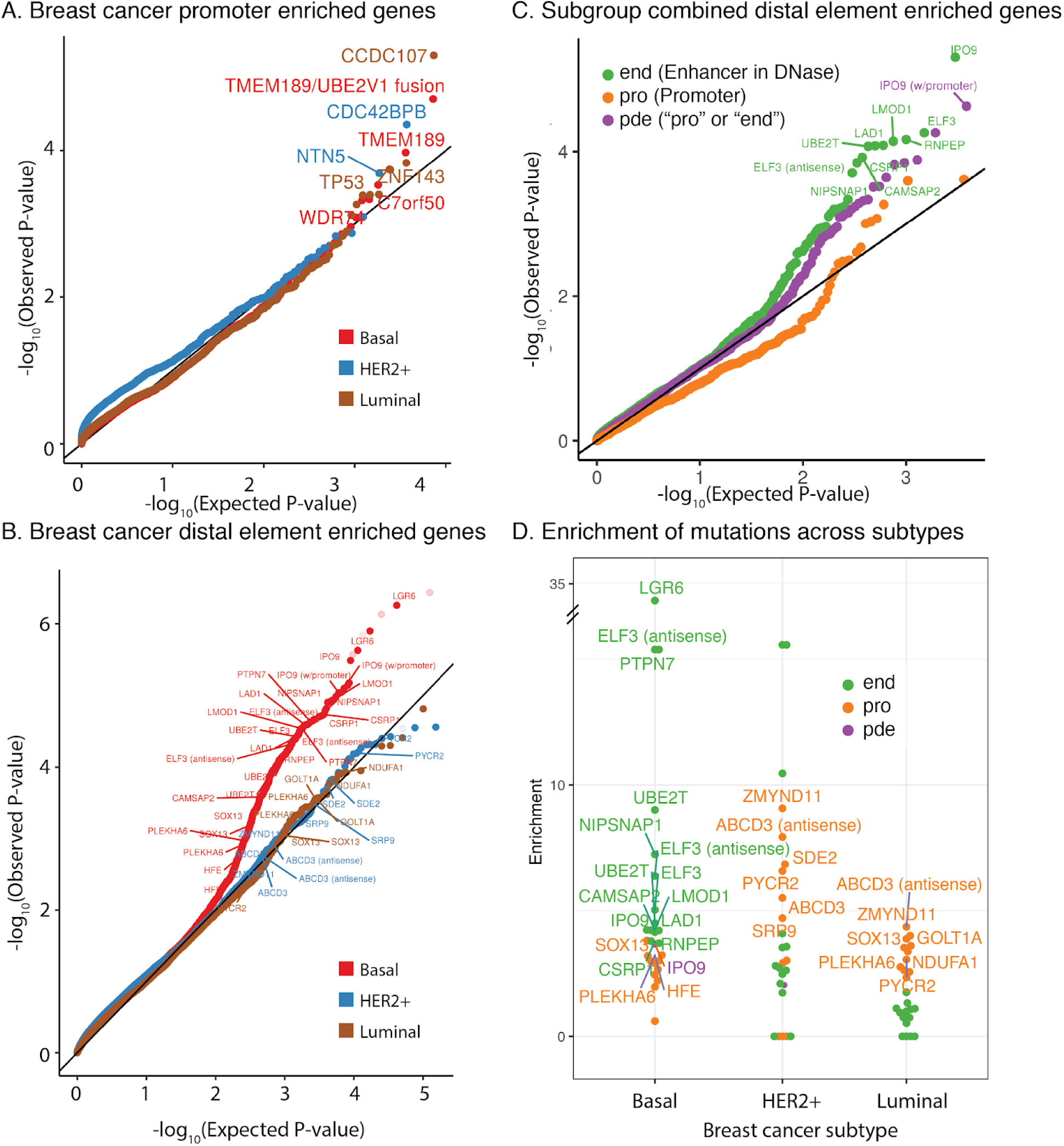
Recurrent regulatory mutations in breast cancer. **A**.**Breast cancer promoter enriched genes**. Quantile-quantile plots of promoter mutations across breast subtypes (Basal, Luminal, HER2+). **B**.**Breast cancer distal element enriched genes**. Quantile-quantile plots of distal regulatory mutations in each breast cancer subtype. **C**.**Subgroup combined distal element enriched genes**. Quantile-quantile plot of different regulatory states, combined across subtypes. Only element-level definitions are shown, either enhancer and DNase (end), promoter or enhancer in DNase (pde), or promoter regardless of DNase status (pro). **D**.**Enrichment of mutations across subtypes**. Enrichment of significantly associated states from the combined analysis. Each dot within a given cancer type represents a single significantly associated gene, and each gene is repeated across all three cohorts to show relative enrichments of associated genes. Only element-level definitions are shown, either enhancer and DNase (end), promoter or DNase enhancer (pde), or promoter regardless of DNase (pro). Note that the promoter chromatin state is frequently observed in highly active enhancer elements as well as promoters themselves.

We further evaluated mutational burden in topological domains from the progenitor human mammary epithelial (HMEC) cells, the closest normal breast cell type with comprehensive epigenomic data (Rao et al. 2014) and observed a significant enrichment in promoter variants for the topological domain containing *PLEKHA6* (Supplemental Figure 2D). The differences between the enhancer-gene linked enrichments and topological domain enrichments is likely because many regulatory regions in a given topological domain do not contribute globally to the expression of genes that reside within that domain (Degner et al., 2012; Gasperini et al., 2019; Kasowski et al., 2013; Kilpinen et al., 2013; McVicker et al., 2013).

### Identification of IPO9 as a putative breast cancer oncogene

We next sought to evaluate whether individuals with mutations in *IPO9* regulatory regions had altered *IPO9* expression. *IPO9* was highly expressed in MCF-7, which contains a mutation in the *IPO9* regulatory region, but not in HMEC cells, consistent with its dysregulation in malignancy. In the independent METABRIC cohort, *IPO9* expression was higher in Basal subtype tumors (Supplemental Figure 3A). Additionally, *IPO9* (1q32) is amplified in 26% of early stage breast cancers in the METABRIC cohort and 22% of advanced breast cancers in the Metastatic Breast Cancer Project (Figure 3A). Among the 560 breast cancer patients with WGS data, only a subset (n=268) had matched RNA-seq data, four of which had *IPO9* mutations. While underpowered to detect an eQTL signal, *IPO9* expression was higher in patients with *IPO9* regulatory mutations (Supplemental Figure 3B). In addition, when examining three validation cohorts of whole genome sequenced tumors, we observed an additional 19 individuals mutated in DNase regions of enhancer-marked chromatin at *IPO9* (Figure 3B). Collectively, these data suggest that increased *IPO9* expression can occur through a variety of mechanisms, including gene amplification, distal regulatory mutations, and proximal mutations at the promoter, consistent with known oncogenes.

**Figure 3:**
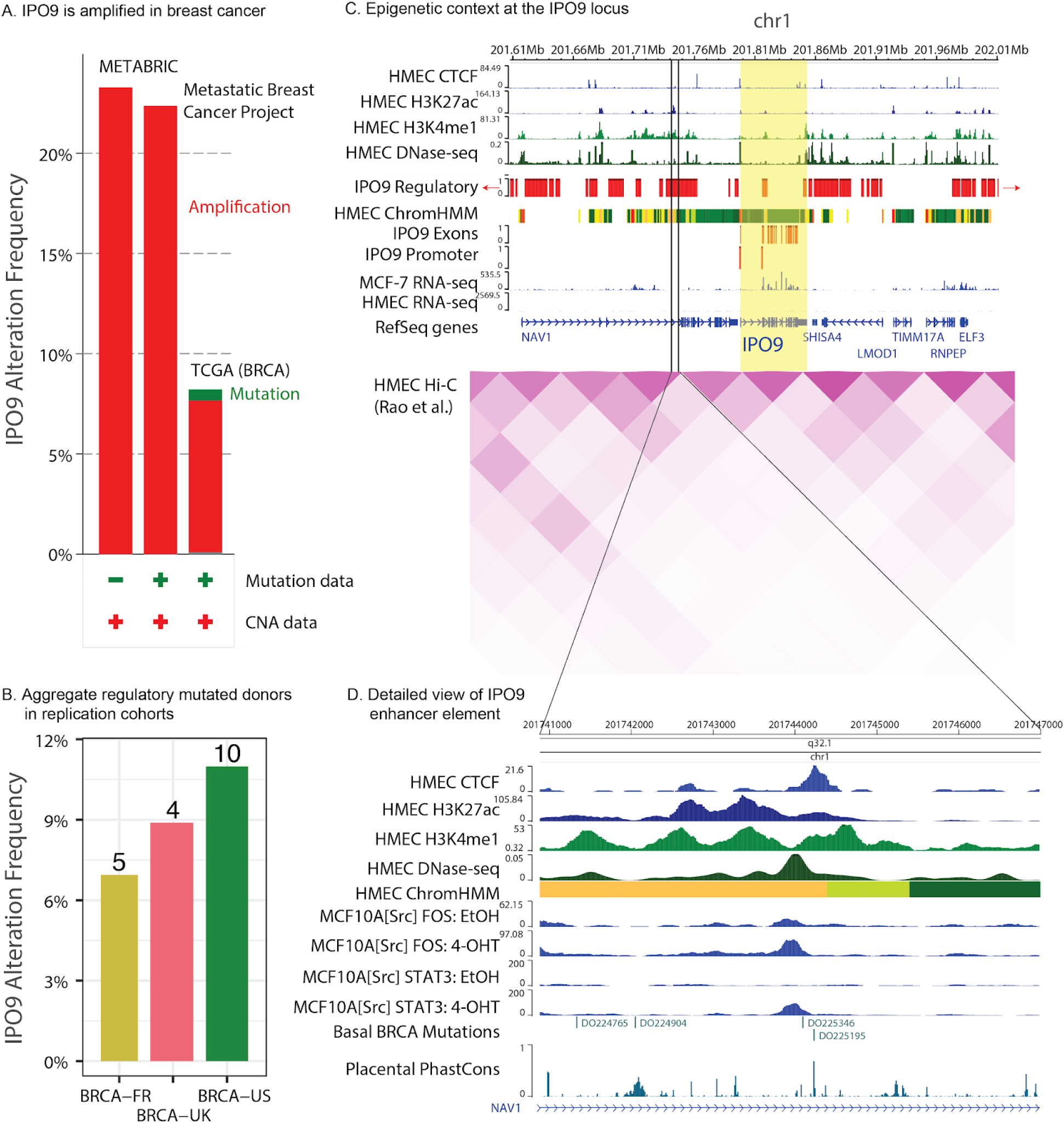
*IPO9* is recurrently altered in breast cancer. ***A***.***IPO9* is amplified in breast cancer**. *IPO9* is frequently amplified in breast cancer across three non-overlapping cohorts: METABRIC (Curtis et al., 2012; Rueda et al., 2019), the Metastatic Breast Cancer Project (Wagle et al., 2016), and The Cancer Genome Atlas (Gao et al., 2013; Liu et al., 2018). There are very few coding mutations in the Metastatic Breast Cancer Project and TCGA. **B**.**Aggregate regulatory mutate donors in replication cohorts**. *IPO9* regulatory region mutations were evaluated in three whole-genome sequenced validation cohorts: BRCA-UK and BRCA-US from PCAWG (ICGC/TCGA Pan-Cancer Analysis of WholeGenomes Consortium, 2020), and BRCA-FR (HER2+ amplified donors) from ICGC (Ferrari et al., 2016). Y axis, fraction of donors with regulatory mutations, with number of mutated donors shown above each bar. All three cohorts show a consistent proportion of donors (∼9%) with mutations in DNase-hypersensitive, enhancer marked regions associated with the *IPO9* promoter. **C**.**Epigenetic context at the *IPO9* locus**. ChIP-seq of histone modifications and CTCF, and open chromatin measured with DNase-seq, in human mammary epithelial cells (HMECs) are shown, as well as the aggregated chromatin state annotations in the two HMEC samples from Roadmap. In addition, the coding and non-coding elements tested for IPO9 are also indicated in red, and the expression of genes in the region is shown for both HMEC cells and MCF-7 breast cancer cells, showing the striking increased expression in MCF-7. Hi-C of HMEC cells (Rao et al., 2014) reveals a domain spanning the majority of regulatory elements (Zhou et al., 2015). **D**.**Detailed view of *IPO9* enhancer elements**. Detailed view of mutational context at an active element in an intron of *NAV1*. The 4-OHT response ChIP-seq profiles in MCF-7 cells and conservation tracks indicates that mutations are primarily located in regions of high activity or conservation.

The epigenetic landscape of breast cancer surrounding the *IPO9* locus is complex and includes large open chromatin regions (defined using DNase-seq), actively transcribed genes (RNA-seq), and regulatory elements (H3K27ac ChIP-seq; Figure 3C). Hi-C data from HMEC cells (Rao et al. 2014) suggests that *IPO9* lies at the boundary of two topological domains, similar to that reported for other regulatory mutations in cancer (Flavahan et al., 2016; Hnisz et al., 2016). We next examined individual regulatory elements containing mutations. One such highly mutated element was located in an intron of *NAV1*, approximately 50Kb away from the *IPO9* promoter and 120Kb away from the *NAV1* promoter (Figure 3D). This element contains a CTCF binding site, active H3K27ac and H3K4me1 marks, as well as a number of conserved regions and DNase hypersensitivity sites. Across all tumors with WGS, there were four breast cancer patients each with a single mutation in this enhancer: one mutation located in a conserved region ∼800bp away, a second located directly adjacent to the CTCF binding site, and two more with mutations located in the DNase hypersensitivity site that is associated with increased STAT3 and FOS binding upon estrogen stimulation in MCF-10A cells (ENCODE Project Consortium, 2012). A similar trend was observed in the *IPO9* UTR, where four regulatory mutations were also present (Supplemental Figure 3C). Together, these data implicate somatic alterations in *IPO9* regulatory elements in breast cancer pathogenesis, as further explored below.

### Pan-cancer aggregate regulatory analysis discovers functional driver genes

We next expanded our analyses to catalogue pan-cancer regulatory driver mutations. We first individually examined the same 20,209 genes used in the breast cancer analysis. As a baseline, when considering all chromatin states rather than restricting to active states, canonical non-coding variants in the *TERT* promoter were observed, as previously reported (Horn et al., 2013; Huang et al., 2013; Vinagre et al., 2013). Enrichment was even stronger when analyses were restricted to active promoters for the cancer type of interest (28-fold versus 14.9-fold enriched). Therefore, for each cancer type we examined the mutational enrichment in the TSS regions using the corresponding active chromatin state information for that type of cancer (Methods). This analysis revealed enrichment in the promoters of the canonical oncogenes *BCL2, TP53, TERT*, and *CXCR4*. We also aggregated the enrichment information across cancer types, which revealed an overlapping, but distinct, set of promoters, including those for *BTG1, CCL15, TERT*, and *TP53* (Supplemental Figure 4C). Thus aggregating promoter mutations across cell types validates canonical driver genes, including *TP53* and *TERT*.

**Figure 4:**
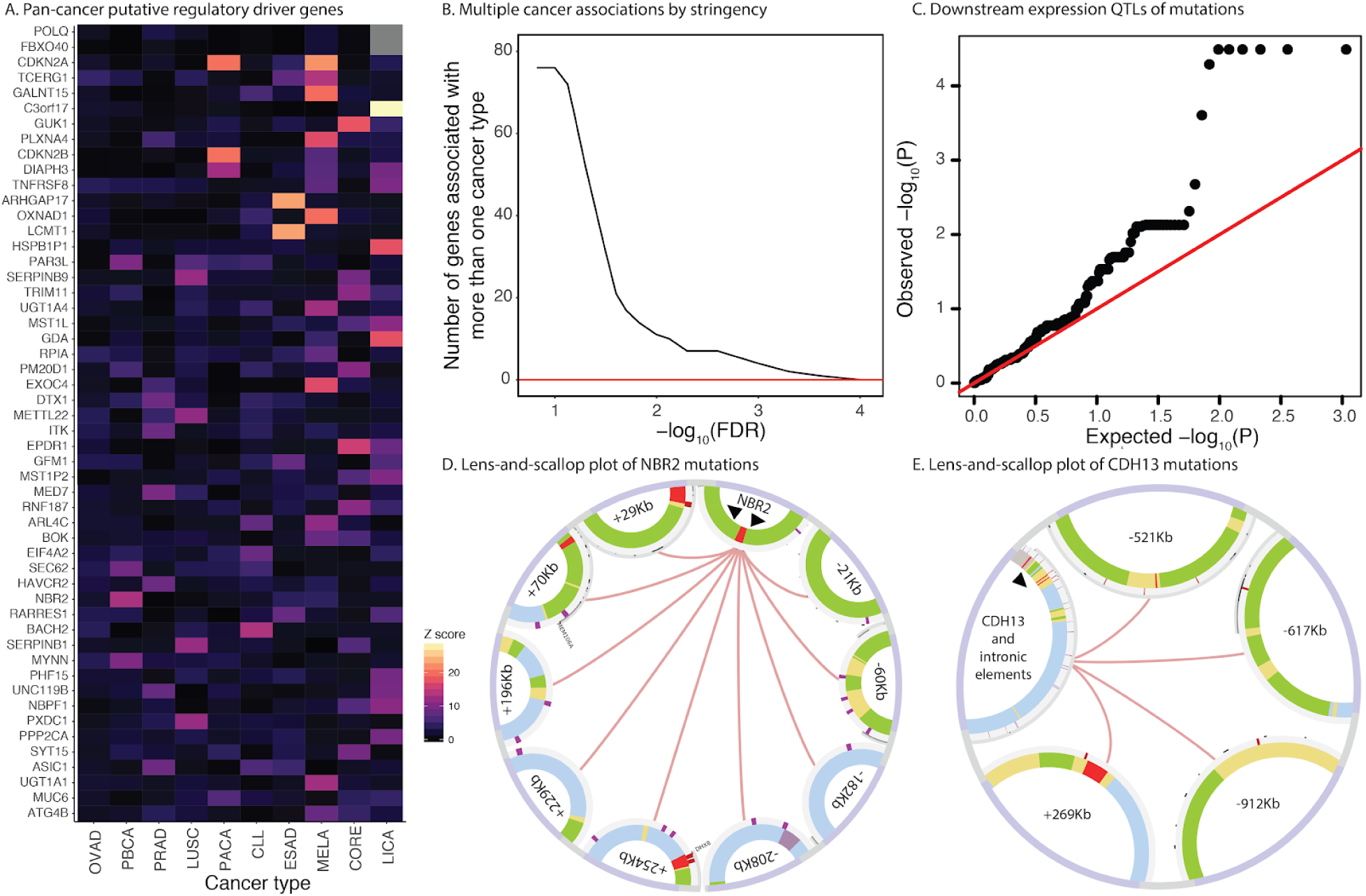
Pan-cancer regulatory mutations have downstream effects on gene expression. **A**.**Pan-cancer putative regulatory driver genes**. The shared landscape of regulatory alterations. Individual cancer types exhibit some uniquely significant genes, whereas other genes are recurrently mutated across cancer types. **B**.**Multiple cancer associations by stringency**. Recurrence of genes across cancer types. Even at increasingly stringent FDR cutoffs, many genes harbor recurrent aggregated regulatory mutations across multiple cancer types. **C**.**Expression QTLs for recurrently mutated regulatory regions**. Overall association of recurrently mutated genes with expression changes. The quantile-quantile plot shows significant changes in expression, as inferred from RNA-seq expression data of mutated versus non-mutated individuals. **D**.**Pan-cancer lens-and-scallop plot of *NBR2* mutations**. Variants are marked with red lines on the outer circle, with regions around mutated regulatory elements shown. Inner circles depict the chromatin state annotations corresponding to the mutated elements. Innermost black arrows at the gene locus mark promoters of *BRCA1* and *NBR2*. **E**.**Pan-cancer lens-and-scallop plot of *CDH13* mutations**. Variants are marked with red lines on the outer circle, with regions around mutated regulatory elements shown. Inner circles depict the chromatin state annotations corresponding to the mutated elements. Intronic elements are shown on gene locus for brevity. Innermost black arrow on gene locus marks promoter of *CDH13*.

We subsequently performed an aggregated distal regulatory element analysis, where we initially employed a parametric approximation (Methods) and then validated significant results with permutation testing. In contrast to methods that focus exclusively on canonical promoter mutations, by aggregated distal regulatory state-specific mutations, we identify numerous novel associations, including both cancer-specific (n = 183) and pan-cancer (n = 40) mutated gene landscapes (Figure 4A, FDR of 10%, Supplemental Tables 5-6). For genes with at least one cancer-specific enrichment, we quantified the significance across more than one cancer type via increasingly stringent FDR cutoffs (Figure 4B).

One example of a hypermutated distal region was a segment associated with *OXNAD1* and *GALNT15*, located 30kb apart. The aggregated distal regions for these genes were specifically overburdened by mutations in CLL and melanoma (enrichment = 4.5 and 1.63-fold, FDR-adjusted q-value = 0.058 and 0.078), and *OXNAD1* was previously reported to be overburdened with promoter mutations in melanoma (Denisova et al., 2015). Additionally, regulatory elements of the non-coding RNA transcript AC090953.1 located within an intron of *GALNT15* was also overburdened with mutations (enrichment = 2.76, q-value = 0.078), though the enhancers overlap substantially with that of *OXNAD1* (Supplemental Table 7). Similar to germline expression QTLs (Tong et al., 2017), co-regulation might mediate this shared enrichment signal. The *TCERG1* gene similarly harbored more mutations (n=27) than expected by chance (n=3.8; q < 0.094) across diverse cancer types, with enrichment in melanoma, esophageal, and ovarian cancers. *TCERG1* is a pro-apoptotic transcriptional elongation factor (Montes et al., 2015) implicated in cancer progression (Bailey et al., 2018; Forbes et al., 2017; Gao et al., 2013) with two mutational hotspots in nearby coding regions of the gene (Supplemental Figure 4G).

We further noted that the distribution of mutations varied significantly between promoter and distal elements for putative drivers. For instance, *OXNAD1* primarily harbored promoter state mutations, whereas *IFI16* and *PYHIN1* share an enhancer element (chr1:158968600-158969600) with mutations in 11 esophageal cancer patients (Supplemental Table 8). Both of these sites would likely be detected with methods that examine individual regulatory elements. However, other genes, such as *BRCA1*/*NBR2* (Figure 4E) and *CDH13* (Figure 4F), were overburdened with variants distributed across multiple elements (e.g. promoters and distal elements), and hence would be overlooked using conventional approaches, including those put forth in recent state-of-the-art single element analyses (Rheinbay et al., 2020).

We further sought to evaluate whether our aggregated non-coding cell-type aware driver discovery method can also recover known pan-cancer drivers of disease in coding regions and UTRs. To this end, we focused on mutations in the “transcribed” chromatin state, corresponding to active genes (Joshi and Struhl, 2005). After removing genes for which the whole gene body lacked H3K36me3, and using Fisher’s method to combine p-values across cancer types, we confirmed the significant enrichment of mutations in known driver genes *TP53, BRAF, NRAS, SMAD4*, and *MUC3* (Supplemental Figure 4C, Supplemental Table 9-11, all but MUC3 reported in Rheinbay et al., 2020). We also observed associations the UTR of *NOTCH1* in CLL (Lobry et al., 2011) (4 patients, 48-fold enriched, q < 0.055), and *AHSA2* and *USP34* in pediatric brain cancers (7 and 6 patients, 20.5-fold and 41-fold enriched, q < 0.0248 and q < 0.0245). Overall, driver genes discovered using a cell type aware model overlapped with those reported previously, but represent only a subset of those discovered using aggregated noncoding elements, highlighting the power of our method to expand the non-coding mutational landscape of cancer.

### Recurrently mutated regulatory regions are associated with cell growth defects

Our regulatory mutation analysis revealed a novel set of genes implicated in cancer. To determine whether these genes are important for cell proliferation, we used genome-wide CRISPR screen data from Project Achilles (Meyers et al., 2017). These analyses indicate that genes enriched for distal mutations tend to be highly deleterious (Figure 5A). Although both distal- and promoter-mutated genes were enriched for deleterious effects (Figure 5B, Supplemental Table 12), knockout of genes with distal regulatory mutations had effects on cell growth comparable to coding mutations. Some genes were essential in nearly all cancer cell lines, including *MED8, GUK1*, and *SDE2* (Figure 5C), whereas others had cancer type specific growth effects (mostly deleterious). For example, *TMEM189* had severe growth defects in leukemia (intercept −0.2 across all lines; leukemia average −0.52, p = 0.038, Supplemental Table 13) and *MAPK1* was less deleterious in myeloma and kidney cell lines (intercept −0.36 across all lines; kidney average 0.044, p = 0.049 and myeloma average 0.12, p = 0.038, Supplemental Table 14). Others were subtype specific - most notable was *PAX5*, where the intercept across cell lines was 0.04 (p = 0.70), but in lymphoid neoplasms, the regression effect was −0.40 (p = 1.8e-18, Supplemental Table 15). In fact, putative drivers were both more primary cancer type specific (Wilcoxon rank-sum test W = 708190, p = 0.037) and had greater dependency scores (median dependency of −0.125 vs −0.06, Wilcoxon rank-sum test W = 754210, p = 0.009) than other genes.

**Figure 5:**
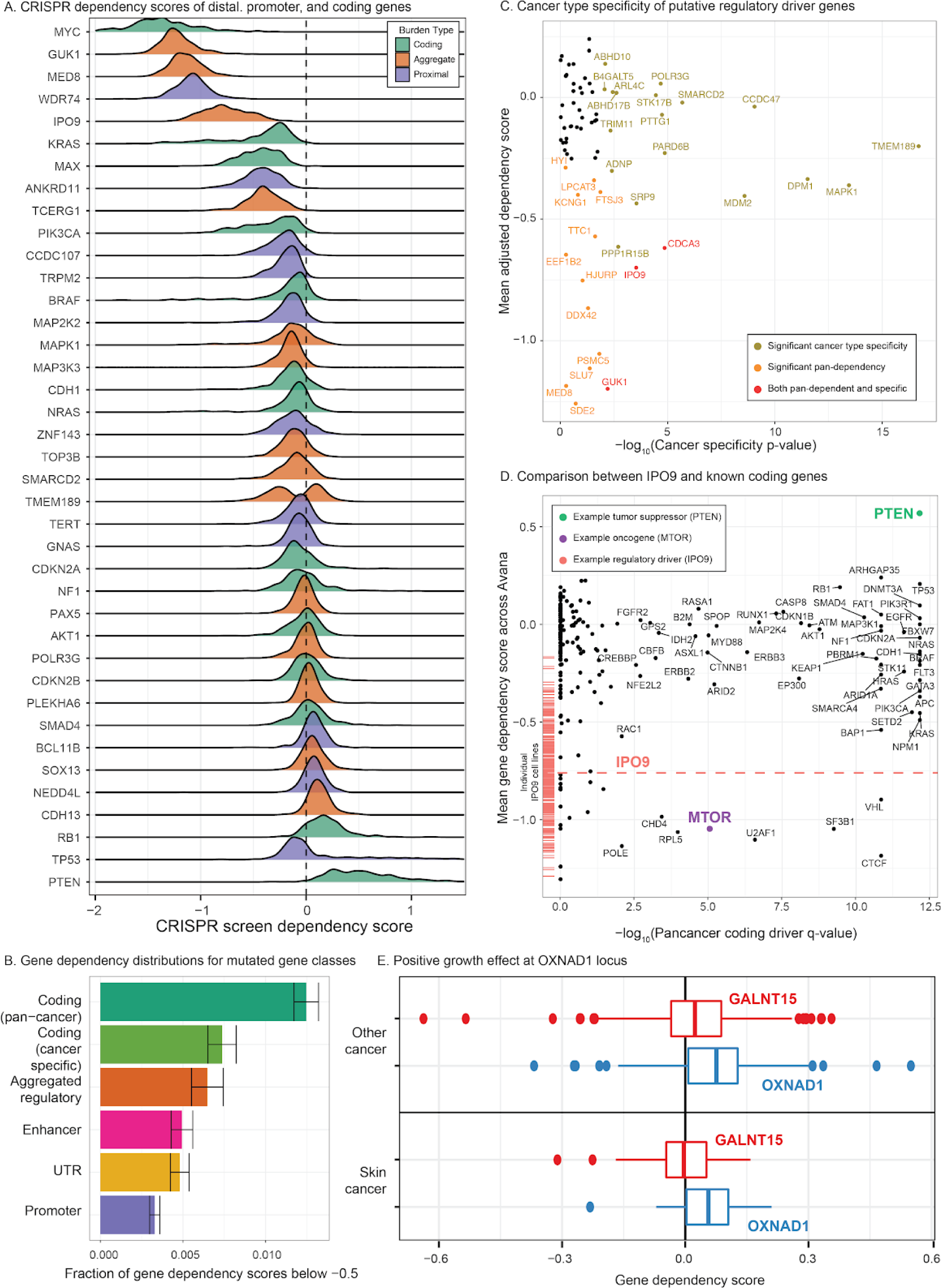
CRISPR screens elucidate distinct mechanisms of regulatory driver function. **A**.**CRISPR dependency scores of distal, promoter, and coding drivers**. Effect on CRISPR growth of known promoters and coding drivers versus novel regulatory drivers. Each distribution is the effects observed across cell lines. Essential gene knockouts have a median dependency score of −1.0, while non-essential gene knockouts have a median dependency score of 0. **B**.**Gene dependency distributions for mutated gene classes**. Across all associated genes, the knockout dependency scores of regulatory, promoter, and coding associated variants relative to the whole genome background. The fraction of gene effects below −0.5 (indicating substantial deleterious effect on proliferation) (Meyers et al., 2017)) are tallied across all genes in the given set. Coding data are from TCGA (Lawrence et al., 2014) and non-aggregate non-coding data are from PCAWG (Rheinbay et al., 2020). **C**.**Cancer type specificity of putative regulatory driver genes**. Each gene was evaluated for cancer type specificity using an F-test (Methods) and the resulting estimates were used to separate genes into those with significant specificity (gold), non-zero aggregate essentiality (orange), both (red), or neither (black). **D**.**Comparison between *IPO9* and known coding genes**. Comparison of coding versus noncoding effects in the Achilles screens. Each dot represents a significant pan- or single-cancer association from Lawrence et al (2014). The red dashed line and bars are the mean estimate and individual estimates of effect for *IPO9*. **E**.**Positive growth effect at *OXNAD1* locus**. Both *GALNT15* and *OXNAD1* have regulatory regions overburdened with mutations, but CRISPR/Cas9 screens reveal a significantly larger positive dependency score for *OXNAD1* compared to *GALNT15* in melanoma and other cell lines.

This suggests that the genes identified through aggregate regulatory mutation analysis have strongly deleterious phenotypic consequences and confer selective advantages through altered gene regulation commensurate with that of coding variants. While strong pan-cancer tumor suppressor genes, such as *PTEN* and *OXNAD1* (newly discovered) (Supplemental Figure 5C), exhibited positive effects on growth, there were very few regulatory genes with positive effects, whereas many genes, such as *IPO9* and the canonical oncogene *MTOR*, showed consistent negative growth effects across all cell lines in Avana (Figure 5D).

### Fine-mapping at the IPO9 locus implicates RNA splicing and processing

*IPO9* knockouts exhibited dramatically reduced proliferation and this gene was pan-essential in both the GeCKO and Avana screens. Indeed, the effect of *IPO9* knockout on proliferation was far larger than other genes in the region (Figure 6A) and persisted across cell types in the independent GeCKO screens (Supplemental Figure 6A). A similar decrease in proliferation was noted for *TIMM17A* in pleural and upper digestive cancers (Supplemental Figure 6B).

**Figure 6:**
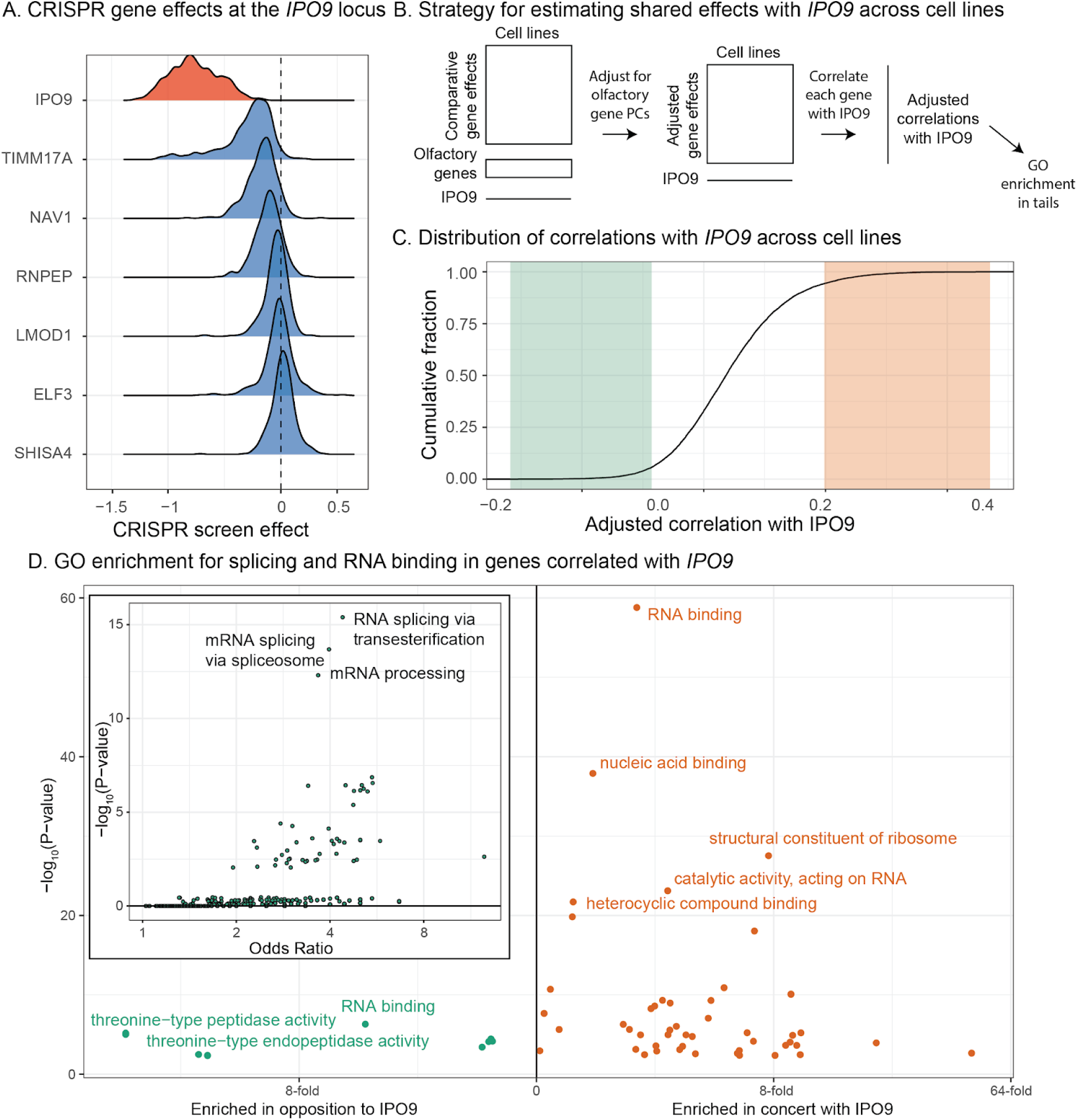
Regulatory and functional characterization of IPO9 using CRISPR screen data. **A**.**CRISPR gene effects at the *IPO9* locus**. Overall distribution of growth effect of *IPO9* versus all other genes at the locus in CRISPR/Cas9 gene knockouts across cancer cell lines (Meyers et al., 2017). **B**.**Strategy for estimating shared effects with *IPO9* across cell lines**. Overall schematic of our method for estimating shared effects across cell lines, similar to previous designs (Boyle et al., 2018). **C**.**Distribution of correlations with *IPO9* across cell lines**. The observed distribution of batch-corrected correlations between *IPO9* and each other genes across Avana and cell lines. Green, negatively correlated genes and orange, positively correlated genes. **D**.**GO enrichment for splicing and RNA binding in genes correlated with *IPO9***. Volcano plots of enrichment for the ranked gene list correlation with *IPO9*, showing a consistent signal of RNA processing. [inset] Volcano plot of enrichment within tail (correlation threshold 0.3), illustrating a substantial enrichment for RNA splicing related genes. Green, negatively correlated genes and orange, positively correlated genes.

This essentiality is further supported by the ExAC database (Lek et al., 2016), where there was a significant depletion of missense variants (z = 3.11) in *IPO9* and the germline probability of loss of function intolerance (pLI) was 1.0. Motivated by this observation, we looked for rare cancer-associated regulatory variants at the locus using the Oxford Brain Imaging Genetics Server (Elliott et al., 2018), and found a variant, rs150641471, in an intron of *NAV1* 50kb from the *IPO9* promoter, which was associated with malignant thyroid neoplasm (OR = 1.05, p = 3e-22), diffuse large cell lymphoma (OR = 1.1, p = 6.6e-12), and leukaemia (OR = 1.004, p = 2.2e-6). This is consistent with transposon screens in mice, which have implicated *IPO9* in hematopoietic malignancy (Guo et al., 2016).

To further characterize the role of *IPO9* in cancer progression, we correlated the gene-level growth effects for *IPO9* with all other genes (Figure 5G-H) following normalization, as previously described (Boyle et al., 2018) (Methods, Supplemental Table 16). Gene ontology (GO) analysis of the 168 genes for which proliferation across cell lines had a correlation greater than 0.3 with *IPO9* revealed the striking enrichment of non-coding RNA metabolic processes (7.29-fold, FDR adjusted q = 7.55e-16, Supplemental Table 6) and catalytic activity on RNA (5.32-fold, q = 1.02e-4). Meanwhile, the most negatively correlated genes include those involved in mRNA splicing via transesterification (4.24-fold enriched in 1000 most negatively correlated genes, q = 1.36e-16; Figure 5I). These results implicate *IPO9* in RNA splicing and processing.

### Recurrently mutated regulatory regions are associated with patient outcomes

Since mutations in regulatory regions often result in gene expression changes, we next examined the association between the expression of genes with recurrently mutated regulatory regions and clinical outcome. We evaluated the specificity of survival associations across 27 cancer types with sufficient clinical information and follow-up duration from the TCGA Pan-Cancer Atlas, the largest compendium of cancer genomes that did not overlap with our non-TCGA ICGC discovery cohort (Bailey et al., 2018; Liu et al., 2018) (Supplemental Figure 6G). In order to limit the number of hypotheses tested, we only evaluated the association between *IPO9, MED8, OXNAD1, PLEKHA6*, and *GUK1* expression and survival. While the trends varied between cancer types, *IPO9* (expression-increasing, risk-increasing), *MED8* (expression-increasing, risk-increasing), and *OXNAD1* (expression-increasing, risk-decreasing) were associated with survival across multiple cancer types (Figure 7A-D, Supplemental Tables 17-18, Supplemental Figure 7F,H-I, after adjusting for key clinical covariates and copy number at that locus, Methods). In addition, increased *PLEKHA6* expression was protective in bladder cancer and lung squamous cell cancer, and risk-increasing in clear cell renal cell cancer.

**Figure 7:**
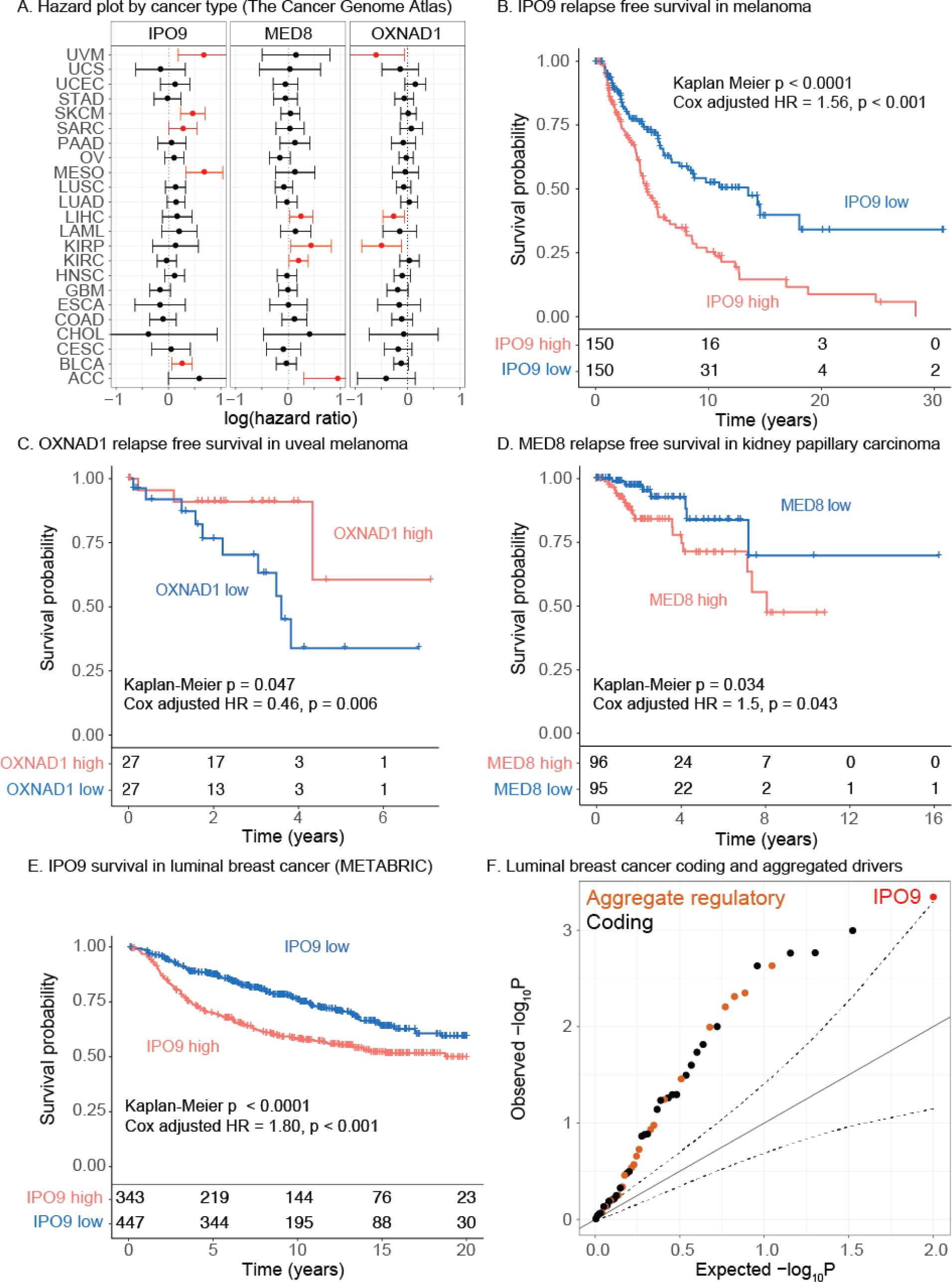
Recurrently mutated genes are associated with clinical outcome. **A**.**Hazard plot by cancer type**. Forest plot for *IPO9, MED8*, and *OXNAD1* from expression and relapse-free survival Cox proportional hazards in TCGA across 23 well-powered cancer types (breast was excluded due to limited followup duration in TCGA). Data for *GUK1* and *PLEKHA6* are reported in Supplemental Figure 6. ***B***.***IPO9* relapse free survival in melanoma (TCGA)**. Kaplan-Meier analysis of the association between *IPO9* expression and relapse free survival in the TCGA melanoma cohort. Cox Proportional Hazards Ratios are also reported. Corresponding forest plot in Supplemental Figure 7, and uncensored counts presented below the axis for each timepoint. ***C***.***OXNAD1* relapse free survival associations in uveal melanoma (TCGA)**. Kaplan-Meier analysis of the association between *OXNAD1* expression and relapse free survival in the TCGA uveal melanoma cohort. Cox Proportional Hazards Ratios are also reported. Corresponding forest plot in Supplemental Figure 7, and uncensored counts presented below the axis for each timepoint. ***D***.***MED8* relapse free survival associations in kidney papillary carcinoma (TCGA)**. Kaplan-Meier analysis of the association between *MED8* expression and relapse free survival in the TCGA ukidney papillary carcinoma cohort. Cox Proportional Hazards Ratios are also reported. Corresponding forest plot in Supplemental Figure 7, and uncensored counts presented below the axis for each timepoint. **E**.**IPO9 relapse free survival associations in luminal breast cancer (METABRIC)**. Kaplan-Meier analysis of the association between *IPO9* expression and relapse free survival in the METABRIC breast cancer cohort. Cox Proportional Hazards Ratios are also reported. Corresponding forest plot and forest plot for all cancers in Supplemental Figure 7, and uncensored counts presented below the axis for each timepoint. **F**.**Luminal breast cancer coding and aggregated drivers**. Quantile-quantile plot of gene expression-survival associations based on disease-free survival in luminal cases in METABRIC. The distribution covers recurrently altered coding variants from TCGA or aggregated regulatory genes from our study (n = 50), revealing enrichment for survival associations. Corresponding plot for all tumors in Supplemental Figure 7.

Next, we sought to evaluate cell-type specific driver effects and their prognostic associations. We initially focused on breast cancer, given the large sample size and long-term clinical follow-up available in the METABRIC cohort (Curtis et al., 2012; Rueda et al., 2019). *IPO9* expression was significantly associated with relapse free survival (RFS) in Kaplan Meier analysis (p < 0.0001) and remained significant in a Cox proportional hazard analysis adjusted for age, tumor grade and size, subtype, and copy number (HR = 1.31 [1.03, 1.7], p = 0.027, Supplemental Figure 7D, Supplemental Tables 19-22) (Methods). We further evaluated this association after stratifying for breast cancer subgroups, revealing an even more striking relationship between IPO9 expression and relapse-free survival in luminal breast cancers (HR = 1.80 [1.29, 2.51], p < 0.001, Figure 7E, Supplemental Figure 7E, Supplemental Tables 23-26).

Encouraged by this result, we evaluated the association between the expression of all genes (n = 50) harboring recurrent regulatory or coding mutations from TCGA and outcome in the METABRIC breast cancer cohort, for. A clear inflation of p-values is noted, suggesting a number of genes are associated with survival. In the METABRIC cohort (Supplemental Figure 7F-H), *IPO9* was the fourth most significant gene, with *SDE2*, which also exhibited large CRISPR growth effects, being the most significant distal association. Of note, *IPO9* expression was most strongly associated with relapse free survival in luminal cases (Figure 7F). The distribution was similar for overall survival, disease specific survival, and distant relapse (Supplemental Figure 7A-C). These findings indicate that genes harboring recurrent regulatory mutations are associated with patient prognosis, cementing their relevance in human cancers.

## Discussion

Here we present a powerful framework to identify non-coding cancer driver genes based on two key principles: aggregation of cell type specific regulatory elements and cell type specific activity to identify novel non-coding driver gene mutations across diverse cancer types. This approach defines driver mutations in multiple regulatory elements simultaneously. Indeed, many regions and associated genes were not identified previously. We demonstrate that mutations in the promoter of *OXNAD1* are likely oncogenic, consistent with previous claims (Denisova et al., 2015). Further, we identify a *IPO9*, a nuclear actin transporter, implicated in mRNA metabolism and alternative splicing, as a putative oncogene in breast cancer, melanoma, bladder cancer, and mesothelioma. In addition to *IPO9*, other newly identified regulatory driver genes, including *SRSF2* and *TCERG1*, also modulate alternative splicing (Koedoot et al., 2019; Montes et al., 2015; Pearson et al., 2008), suggesting a shared functional basis for these enrichments, similar to that also seen for alternative splicing in coding mutations (Watson et al., 2013).

Previous work has implicated *IPO9* in nuclear actin remodeling and adherence of keratinocytes (Sharili et al., 2016), as well as in transcriptional control (Dopie et al., 2012) and interferon signaling (Matsumiya et al., 2013). More recently, nuclear actin has been implicated in the transport of homologous recombination double stranded breaks to the periphery, where they can be efficiently repaired (Caridi et al., 2018). In addition, nuclear actin dynamics, mediated by *IPO9* and *XPO6*, have the potential to modulate mRNA splicing through disruption of *SMN2 (Viita et al., 2019*). Alternative splicing and other co-transcriptional metabolic processes acting on RNA are important for cancer development (David and Manley, 2010; Koedoot et al., 2019), suggesting a multitude of direct targets in promoting the hallmarks of cancer (Hanahan and Weinberg, 2011). These diverse roles of nuclear actin in cellular proliferation and transcription are consistent with our findings of mutational enrichment in *IPO9* regulatory regions and the association between elevated expression of *IPO9* and shorter relapse-free and overall survival in multiple cancer types. Together, this motivates further investigation of the mechanism and diversity of nuclear actin as a class of oncogenes using high-content imaging platforms with drug libraries and/or CRISPR tools.

More broadly, our method has uncovered a unique set of recurrently mutated genes not identified through conventional means, including recent large-scale non-coding analyses (Rheinbay et al., 2020). The observation that aggregated regulatory signals harbor enrichment not evident from the analysis of individual elements is reminiscent of progress in exome testing. Initial studies first evaluated individual coding variants, and later found increased power in gene-level burden tests. This suggests that applying novel approaches to the analysis of non-coding regions, including the development of specific driver detection tools, is of value.

The strong growth phenotypes of these genes identified via CRISPR/Cas9 screens suggests that they might be constrained for coding variation, and that distal regulatory elements with slight expression-altering mutations might jointly control expression at multiple loci, akin to polygenic models in genome-wide association studies. These findings also highlight the power of large scale genetic screens to inform driver gene discovery and we identify an excess number of mutated genes with large deleterious growth effects. It is worth noting, however, that loss of large-effect tumor suppressors during serial passaging is anticipated, and such genes would not be identified in this analysis. Finally, we illustrate how loss of function genetic screens can be used to fine map causal genes, evaluate cancer type specificity and determine functional mechanisms, including direct annotation of pathways.

In sum, we present a general approach to identify regulatory regions enriched for mutations while simultaneously correcting for background mutation rates. The application of this approach to WGS data from 11 cancer types, lead to the identification of multiple novel non-coding driver genes, supported by orthogonal validation of their pan-cancer growth effects and prognostic associations. Of note, these findings likely represent just the beginning, and we anticipate that additional non-coding drivers will be identified through the application of this new cell-type aware, analytic framework to the increasing number of WGS cancer datasets being generated with implications for personal genome interpretation and prognosis. Together, we believe that improved methods like these, as well as additional genomic and other omics data, will begin a new large-scale effort to discover and interrogate regulatory drivers in cancer.

## Acknowledgements

N.S.-A. is supported by the Department of Defense through a National Defense Science and Engineering Grant and by a Stanford Graduate Fellowship. This work was supported by grants from the NIH ENCODE/Production Center for Mapping Regulatory Regions of the Human Genome - 3U54-HG006996-04S1 (M.S.), CEGS/Center for Personal Dynamic Regulomes - 2RM1HG007735-06 (M.S., C.C.), the NIH Director’s Pioneer Award - DP1-CA238296 (C.C.) and the NCI Cancer Target Discovery and Development Network - U01 CA217851 (C. C.). This work made use of resources provided by the Genetic and Bioinformatic Service Center at Stanford University School of Medicine. We thank C. Suarez for feedback and tumor epigenome matching and M. Wainberg, A. Sanghi, D. Rothschild, C. Smith, H. Ollila, J. Reuter, J. Gruber, M. Pryzbilla and other members of the Curtis, Pritchard and Snyder labs for feedback on the manuscript. This study makes use of patient data from the ICGC, TCGA, and METABRIC studies, as well as epigenetic data from the ENCODE and Roadmap Epigenomics Project and CRISPR screen data from DepMap. We thank all the individuals who contributed to the creation of these resources, and particularly the patients whose biological samples form the basis of this work. This study was performed on the traditional and unceded lands of the Muwekma Ohlone people, and we are grateful for the opportunity to live and work here.

## Author contributions

Conceptualization: N.S.-A. and R.S.

Methodology: N.S.-A., R.S., C.C., and M.P.S.

Software, Analysis, and Validation: N.S.-A. and J.A.S.

Investigation: N.S.-A., J.A.S., C.C. and M.P.S.

Writing and Editing: N.S.-A., J.K.P., C.C. and M.P.S.

Supervision: C.C. and M.P.S.

## Competing interests

C.C. is a scientific advisor to GRAIL and reports stock options, as well as consulting for GRAIL and Genentech. M.P.S. is a co-founder and SAB member of Personalis.

## Data & Code Availability

This study makes use of patient data from the ICGC, TCGA, and METABRIC studies, as well as epigenetic data from the ENCODE and Roadmap Epigenomics Project. Data from TCGA are available publicly through the PanCan Atlas portal (https://gdc.cancer.gov/about-data/publications/pancanatlas) and via application to dbGaP accession phs000178.v1.p1. Data from ICGC are available on the ICGC website (http://icgc.org/). Data from ENCODE and Roadmap are available on the ENCODE website (http://encodeproject.org). Data from DepMap are available on the DepMap website (https://depmap.org/portal/download/). Data from METABRIC are available at the European Genotype-Phenotype Archive under Accession number EGAS00000000083 and as supplementary tables in the current publication (Rueda et al., 2019).

The code needed to implement the methods described in this paper will be published along with the accepted manuscript.

## Supplemental Tables

Supplemental Table 1: **Tumor samples included in the discovery cohort**. The list of all tumors used for initial discovery of driver mutations, including the aggregated tumor type used for these analyses, the original cohort from ICGC, and the donor ID. Cancer type, cohort name, and donor ID are listed.

Supplemental Table 2: **Chromatin state definitions**. The abbreviated names, equation (used internally for specifying the definition), chromatin states, and DNase status of aggregated active chromatin used for the analysis.

Supplemental Table 3: **BRCA combined putative driver list**. List of all putative driver genes discovered in breast cancer using the fisher-combined p-values across cohorts, including the chromatin state tested; resolution of tile resampling employed; mutation rate window; set of chromatin loops evaluated; and expected mutation count across permutations, number of observed mutations, and likewise for number of patients mutated, as well as the empirical value and FDR-adjusted q-value. Only genes with a patient q-value < 0.1 are reported.

Supplemental Table 4: **BRCA combined active promoter and all promoter genes**. List of all genes putatively enriched in promoter mutations, either including all chromatin states or only promoter chromatin annotations (active).

Supplemental Table 5: **Single-cancer coding driver genes**. List of all genes putatively enriched in coding mutations in each single cohort. Mutation, number of mutations observed; patient, number of patients with mutations; permutations, number of permutations run to evaluate significance; mean_mutation, average number of mutations in permutations; mean_patient, average number of patients mutated in permutations; gtmutation, number of permutations with mutation count exceeding the observed; gtpatient, number of permutations with patient count exceeding the observed; p.pt, empirical p-value of patient mutations; q.pt empirical FDR-adjusted p-value of patient mutations.

Supplemental Table 6: **Pan-cancer combined coding drivers**. List of all putative coding genes discovered in the pan-cancer analysis using the fisher-combined p-values across cohorts. FDR cutoff of 10% was used to report genes, and each gene was assessed using the permutation-based approach.

Supplemental Table 7: **Pan-cancer combined coding active drivers**. List of all putative coding genes discovered in the pan-cancer analysis using the fisher-combined p-values across cohorts, but only using mutations located in actively transcribed regions. An FDR cutoff of 10% was used to report genes, and each gene was assessed using the permutation-based approach.

Supplemental Table 8: **Parametric single-cancer putative drivers**. List of all putative single-cancer aggregate regulatory drivers discovered using the parametric models. Cancer, cancer type; links, regulatory element links used; state, chromatin state tested; rmr, window size (bp) for calculating regional mutation rate; mutated, number of mutations observed, mean, number of mutations expected; z, z-score based test statistic; log10pois, log of the p-value for the poisson test; log10chi, log of the p-value for the chi squared test; log10z, log of the test statistic for the Z test; qchi, FDR-adjusted q-value for the chi square test; qpois, FDR-adjusted value for the poisson test; qz, FDR-adjusted q-value for the z test.

Supplemental Table 9: **Pan-cancer combined putative drivers**. List of all putative driver genes discovered in the pan-cancer analysis using the fisher-combined permutation p-values across cohorts. Only genes that were validated with the permutation-based approach are reported. State, chromatin state tested; mutations, number of observed mutations; patients, number of mutated patients. QC is marked “FAIL” for histone, immunoglobulin, and RNA genes excluded from downstream analysis.

Supplemental Table 10: **OXNAD1/GALNT15 MELA mutated elements**. List of mutations from the linked regulatory regions of OXNAD1, GALNT15, and the nearby non-coding RNA. Each row represents a mutation-gene combination, with the corresponding chromatin state and regulatory region annotated.

Supplemental Table 11: **Mutations in a PYHIN1-IFI16 shared enhancer**. List of individual mutations located in the enhancer element shared by PYHIN1 and IFI16 across the esophageal cancer cohort.

Supplemental Table 12: **Essentiality comparison across genes**. The fraction of gene effects labeled essential for genes associated with coding mutations from TCGA (Bailey et al., 2018); coding, promoter, enhancer, or UTR mutations from PCAWG (Rheinbay et al., 2020); and aggregated regulatory regions in either breast cancer or the pan-cancer cohort (this study).

Supplemental Table 13: **Cancer type specificity of TMEM189**. Regression specification for the cancer type specificity of TMEM189, adjusted for olfactory gene essentiality principal components 1-5; gender; and source.

Supplemental Table 14: **Cancer type specificity of MAPK1**. Regression specification for the cancer type specificity of MAPK1, adjusted for olfactory gene essentiality principal components 1-5; gender; and source.

Supplemental Table 15: **Cancer subtype specificity of PAX5**. Regression specification for the cancer type specificity of PAX5, adjusted for olfactory gene essentiality principal components 1-5; cancer type; gender; and source.

Supplemental Table 16: **Essentiality correlation with IPO9**. Table of pairwise batch-corrected correlations between each of the genes evaluated in the Avana screen and IPO9 across all 485 cell lines in the Avana dataset.

Supplemental Table 17: **TCGA per cancer type hazard ratios**. Across each of the 33 cancer types in the PanCanAtlas, the hazard ratio of expression changes for each of the five genes we selected for downstream analysis (*IPO9, PLEKHA6, GUK1, MED8*, and *OXNAD1*).

Supplemental Table 18: **TCGA combined hazard ratios across cancer types**. Combined hazard ratio for the five genes evaluated in multiple cancer types with adequate sample size.

Supplemental Table 19: **Overall survival hazard ratios in METABRIC**. Hazard ratios, for each putative breast cancer driver gene, of expression against overall survival when adjusted for standard clinical covariates.

Supplemental Table 20: **Disease specific survival hazard ratios in METABRIC**. Hazard ratios, for each putative breast cancer driver gene, of expression against disease specific survival when adjusted for standard clinical covariates.

Supplemental Table 21: **Relapse free survival hazard ratios in METABRIC**. Hazard ratios, for each putative breast cancer driver gene, of expression against relapse free survival when adjusted for standard clinical covariates.

Supplemental Table 22: **Disease and relapse free survival hazard ratios in METABRIC**. Hazard ratios, for each putative breast cancer driver gene, of expression against disease- and relapse-free survival when adjusted for standard clinical covariates.

Supplemental Table 23: **Overall survival hazard ratios in METABRIC, luminal cases only**. Hazard ratios, for each putative breast cancer driver gene, of expression against overall survival when adjusted for standard clinical covariates, among luminal cases only.

Supplemental Table 24: **Disease specific survival hazard ratios in METABRIC, luminal cases only**. Hazard ratios, for each putative breast cancer driver gene, of expression against disease specific survival when adjusted for standard clinical covariates, among luminal cases only.

Supplemental Table 25: **Relapse free survival hazard ratios in METABRIC, luminal cases only**. Hazard ratios, for each putative breast cancer driver gene, of expression against relapse free survival when adjusted for standard clinical covariates, among luminal cases only.

Supplemental Table 26: **Disease and relapse free survival hazard ratios in METABRIC, luminal cases only**. Hazard ratios, for each putative breast cancer driver gene, of expression against disease- and relapse-free survival when adjusted for standard clinical covariates, among luminal cases only.

## Methods

### Variant calls and sample inclusion

Tumor types with whole genome sequencing as part of the International Cancer Genome Consortium for which a minimum of 90 individuals were profiled and for whom matched epigenomic data was available from the ENCODE and RoadMap Epigenome projects were selected for inclusion. Germline filtered somatic mutational calls based on whole genome sequencing were used for downstream analyses where individuals with fewer than 100 somatic mutations were excluded (due to limitations in defining chromatin-state-specific mutational effects). Each cancer type was treated as a single cohort, with the exception of breast cancer (BRCA) where additional stratified analyses were performed according to major subgroups (Luminal, ERBB2/Her2-positive, and triple negative breast cancers (TNBC)). The full list of ICGC donor IDs and cohorts is included in Supplemental Table 1. A total of 2634 individuals were included across all cancer types.

METABRIC expression, CNA, clinical, and survival data were downloaded from European Genome-Phenome Archive (EGA). Data from The Cancer Genome Atlas were utilized for expression-survival validation (Liu et al., 2018) and CRISPR analyses (Bailey et al., 2018) and PCAWG was used for CRISPR analyses (Rheinbay et al., 2020).

### Defining chromatin state and open chromatin regions

Chromatin state annotations for all cancer types except prostate were downloaded from the Roadmap Epigenomics Project integrated analyses while DNase hypersensitivity peaks for all cancer types except prostate were downloaded from the ENCODE portal. For prostate cancer, annotations were obtained from GEO:GSE63094 and quantized to chromatin states in 100bp windows using ChromHMM, and used as annotation sources as described previously (Sallari et al., 2017).

We used a stringent filtering step based on sequence uniqueness to avoid miscalling of chromatin states. In brief, three filters were combined to eliminate regions that might have artifactual annotations or missing genotype calls as a result of mappability bias. First, the ENCODE blacklist regions and UCSC hg19 genome assembly gaps were merged together, followed by looking in umap (ENCODE Project Consortium, 2012) and removing non-uniquely-mappable regions. This results in approximately one third of the genome (mostly centromeric and telomeric regions) being masked of repetitive regions.

### Regional mutation rate estimation and null model mutation distribution

While replication timing data are available in some relevant cell types through ENCODE, the vast majority of cancer types have no annotations available. As such, the regional mutation rate was used as an estimate of replication timing, given their high correlation and reproducible effects on mutational spectrum (Stamatoyannopoulos et al., 2009). Two distinct windows of mutation counts were used -- 25kb and 250kb -- and the counts were summed across patients normalized by patient count (so that rates are comparable between cancer types), total number of mutations in the patient, and the window size (to achieve comparable distributions for both 25kb and 250kb windows).

At every nucleotide in the genome, on a per-cancer-type basis, covariates were estimated as the chromatin state (reduced to 7 states: promoter, enhancer, transcribed, repressed, bivalent, heterochromatin, and quiescent), DNase hypersensitivity peaks, and estimated regional mutation rate, the calculation of which is described above.

To ensure the robustness of results, all models were repeated with multiple regional mutation rate windows and nucleotide fragment sizes. For the single nucleotide model, we ran models corrected for stranded trinucleotide context (Alexandrov et al., 2013). Using these distributions, we tested for the enrichment of mutations across active chromatin states. We focused on active regulatory regions as these have previously been implicated in cancer development (Sabarinathan et al., 2016), and because epigenetic alterations in the cell of origin are thought to potentiate cancer development via loss of tumor suppression (Garinis et al., 2002).

### Mapping regulatory elements to genes

Regulatory elements were mapped to genes using Hi-C links, described above, as well as with correlation-based links (Rheinbay et al., 2020) that utilize modules of co-activated enhancers and co-expressed genes across the Roadmap RNA-seq profiled samples. In addition, the core promoter region was added to the tests as relevant, using annotations from the FANTOM5 consortium (FANTOM Consortium and the RIKEN PMI and CLST (DGT) et al., 2014). Histone I genes, immunoglobulin genes, HLA genes, non-coding “AC” genes, and RNA genes were excluded from further analyses due to either their repetitive structure or lack of adequate annotation coverage, respectively.

Promoter elements (n = 57,534) were defined based on the FANTOM5 consortium CAGE sequencing (FANTOM Consortium and the RIKEN PMI and CLST (DGT) et al., 2014). Promoter BED region defintions were then aggregated within each protein coding gene and intersected with chromatin state annotations. Any elements overlapping with collapsed promoter/strong enhancer (Tss or TssFlnk) chromatin states were labeled as active promoters in downstream analysis.

### Estimation of mutational overburdening

Four tests were employed to estimate the overburdening of mutations. In the first approach, a resampling strategy replaced each tile (a region of consecutive bases, between 1bp and 100bp) in the aggregate regulatory landscape with one that has the same reference nucleotide context, regional mutation rate, chromatin state, and open chromatin level. Then the number of mutations is assessed and the significance is calculated through the empirical p-value relative to the genomic background null distribution. This is exact and gives uninflated quantile-quantile plots, but is computationally intensive to calculate, and thus all associations were first run using the parametric models described below, and marginally significant associations were replicated using the permutation test as a final filter. For evaluation of coding gene effects, q-values for enrichment of putative cancer-mutated genes (Lawrence et al., 2014) were downloaded and ordered by their pan-cancer q-value.

As a pre-filter for the pan-cancer runs, where non-parametric tests are prohibitively time consuming, a poisson distribution is used, where the lambda parameter is estimated from the genome-wide distribution of nucleotides that share the same covariates (regional mutation rate, patient, chromatin state, and DNase sensitivity). Every nucleotide is assumed to be independent and the product of the observed values is the overall expectation.

In order to capture putative enriched genes which violate the poisson assumption, a z-score test is used, where the mean mutation count was derived using the same covariates as the Poisson test. Finally, the Cochran-Mantel-Haentzel (CMH) test was used in which chromatin state strata are simultaneously tested for having mutations at an odds ratio other than one. Together, these three tests act as filters to identify only the gene-state-cancer type combinations most likely to be enriched, and those combinations can be further refined using the non-parametric models.

For the non-parametric models, genomic windows of size 1bp, 10bp, or 100bp were stratified by canonical chromatin states and the presence of open chromatin, and within each, normalized regional mutation rate (mutations per megabase per thousand donors) and reference trinucleotide context were recorded. To evaluate a gene, the associated regulatory regions were divided into chromatin states, and the number of tiles of a given size and parameters were tallied. Then, for each permutation, random matched regions were regenerated and tallied from covariate-matched regions of the same length across the genome and summed across the regulatory landscape.

Fisher’s method was used to combine p-values across cancer types. Under this model, we assume that the estimates from the cancer types are independent given the lack of individual-level overlap between studies of different cancer types.

#### Bootstrap validation of mutation enrichment

A validation of the mutation selection process was performed for the Breast cancer association at *IPO9*. Individuals were resampled uniformly at random in the Basal breast cancer subtype and the observed and expected number of mutations were recalculated. Resampling was performed 20 times and the enrichment in both mutation counts (Supplemental Figure 2A) and patient counts (Supplemental Figure 2B) were tallied.

### Survival analyses

For the METABRIC cohort, clinical data, including relapse free survival was obtained from (Rueda et al. Nature 2019), and expression and copy number from EGA. Expression of *IPO9* was adjusted by copy number by regressing the copy number value from the expression. Kaplan-Meier plots were generated with the package “survminer”, where the top 1/3 and bottom 1/3 expression values for each gene were defined as high versus low, respectively. Cox Proportional Hazards Models were generated using the CoxPH function in the survival package, adjusting for relevant clinical covariates, including age, stage, grade, size, number of lymph nodes positive, estrogen and progesterone receptor status, as well as HER2/ERBB2 status. Estrogen receptor (ER) status was not included in the model for luminal tumors since most are ER-positive. For the TCGA outcome analysis, clinical data (overall survival) was obtained from (Liu et al. Cell 2018), and expression (FPKM, upper quantile) and copy number data from gdc.cancer.gov. Expression was log2 transformed and scale normalized. Cox Proportional Hazards Models were generated similar to that for the METABRIC cohort, again adjusting for clinical covariates (when available) including age, stage, gender and grade. Only tumor types with sufficient numbers and follow-up times were used for the main analyses (Liu et al., 2018).

### CRISPR screen and essentiality analyses

CNA-normalized gene effect scores were downloaded from DepMap for the Avana and GeCKO genome wide CRISPR-KO screens (Meyers et al., 2017). These values represent the normalized effect on cell growth for knockout of the given gene, such that negative values are associated with more lethal knockout. However, potential batch effects are present in the reported essentiality scores (Boyle et al., 2018), and we sought to adjust for these in our aggregated analyses. In brief, for the co-essentiality testing with *IPO9* and driver gene list analysis, the whole gene effect score matrix was normalized using a strategy to remove batch effects (Boyle et al., 2018). The matrix was subset to olfactory receptor genes and PCA was performed, followed by removal of the top five principal components of the olfactory receptor gene matrix from the essentiality of every gene. Driver genes from aggregated elements were subset to those with at least three patients mutated and FDR < 20%. For the correlation analysis with *IPO9*, genes were ordered according to observed correlation coefficients across cell lines (using a cutoff of 0.3).

