## Supplemental Figures and Figure Legends for "Convergent mutations in tissue-specific regulatory regions reveal novel cancer drivers"

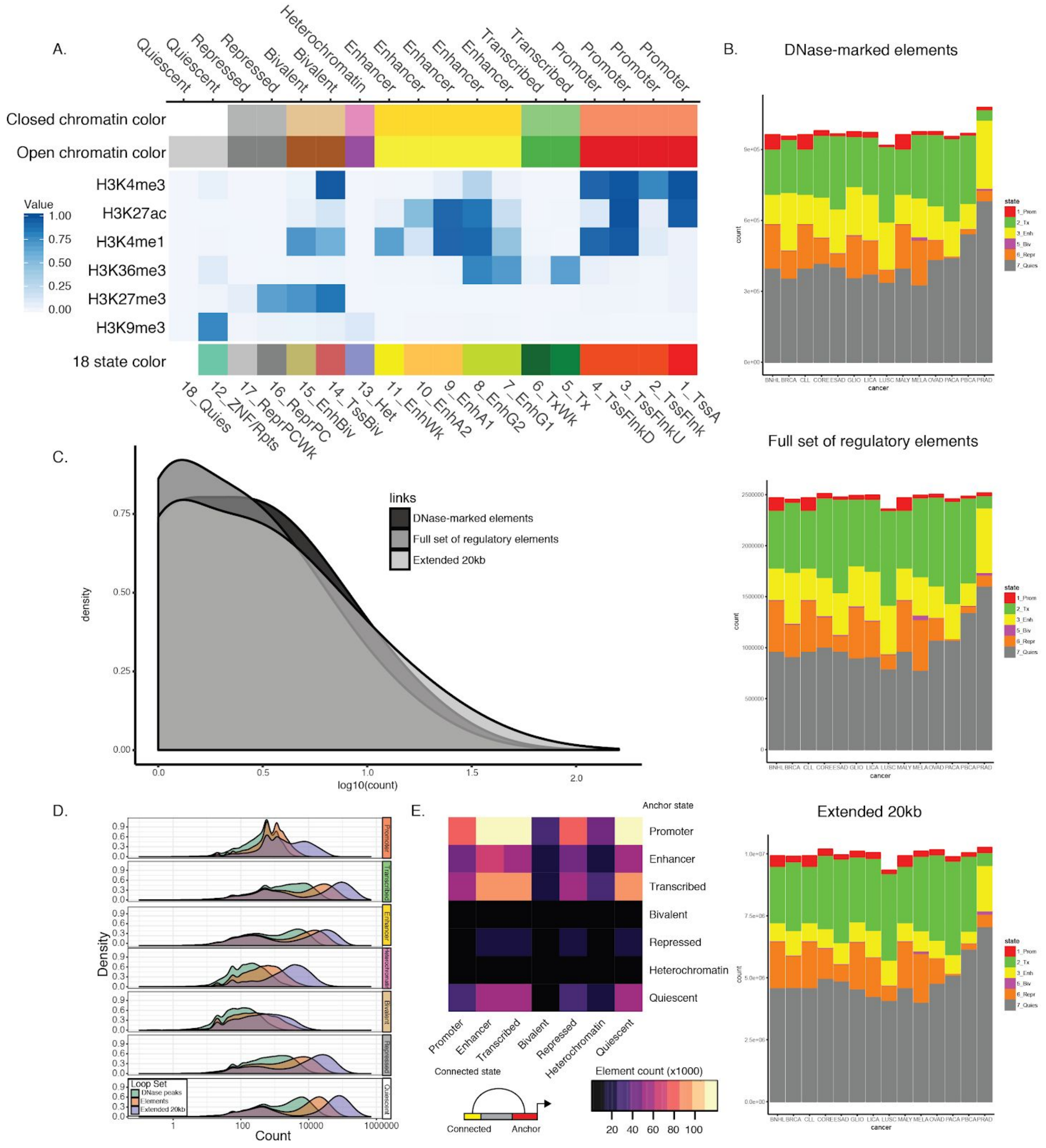


**Supplemental Figure 1: Details of chromatin states and enhancer-gene linking.**

A. **Chromatin state emission parameters.** Chromatin state annotations based on the Roadmap Epigenomics project (Roadmap Epigenomics Consortium et al., 2015). For each chromatin state, the emission parameters in the hidden markov model defining that state for all utilized histone modifications are noted. Reproduced from the Roadmap Epigenomics Project.

B. **Chromatin state distribution across different enhancer linking sets.** Counts of chromatin states across cancer types for elements containing a DNaseI hypersensitive site; at all regulatory elements; and at regulatory elements extended by 20kb in each direction. PRAD is derived from a different study than the other cancer types, resulting in a different distribution (Methods).
C. **Number of genes per enhancer region.** Distribution of the number of genes linked to each enhancer element.
D. **Distribution of enhancer-gene links.** The density of base pairs included in regulatory regions across genes. The three different sets of linked elements represent different stringencies of association; DNase elements are those with strong open chromatin signatures within associated regulatory regions, Elements are those elements themselves, and Extended elements are the elements and 20kb on either side, to help capture regulatory regions which might not have strong direct contact but contact with a gene's promoter through proximity.
E. **Pairwise contact between promoters and enhancers.** The x axis indicates the chromatin state at a given DNase-associated regulatory region, while the y axis indicates the chromatin state at the corresponding promoter. There is a strong divide between regulatory regions and promoters based on "active" regulatory marks. In addition, as expected, there is more enrichment for enhancer-promoter and promoter-enhancer pairs than for either state with itself.

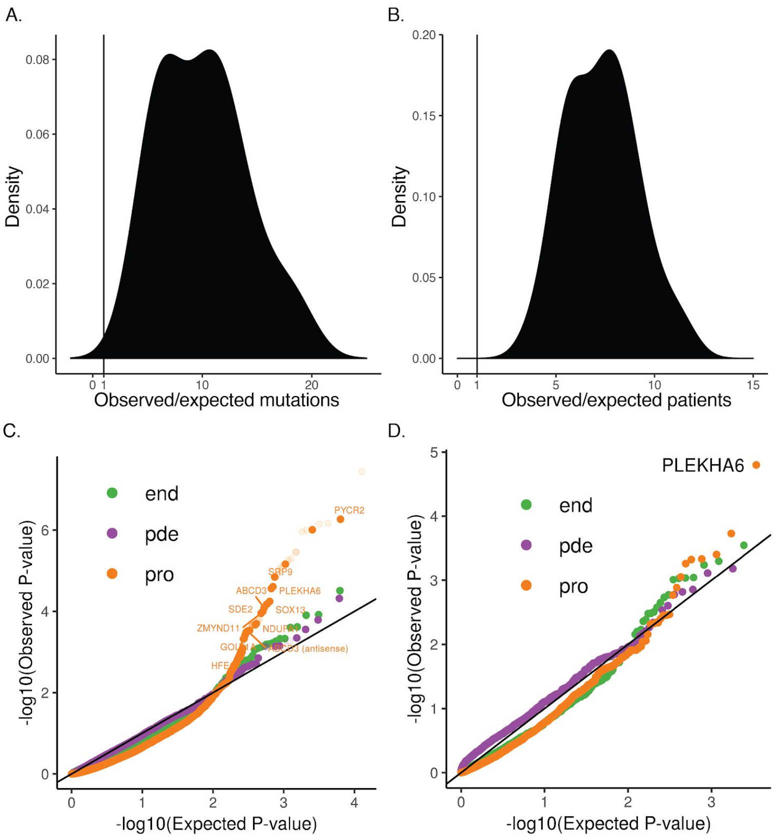


**Supplemental Figure 2: Bootstrap validation of IPO9 mutational landscape.**

A. **Bootstrap validation of *IPO9* mutations.** Enrichment of observed mutations at the IPO9 locus over the expected background for each of 20 iterations (Methods) reveals consistent enrichment over expectation. Bootstraps were performed over individual tumors included in the analysis.
B. **Bootstrap validation of *IPO9* patients mutated.** Enrichment of observed number of patients harboring mutations at the *IPO9* locus over the expected background for each of 20 iterations (Methods) reveals consistent enrichment over expectation. Bootstraps were performed over individual tumors included in the analysis.
C. **Quantile-quantile plot of 20kb-extended regulatory elements.** Observed versus expected p-values when using all regulatory elements and the 20kb surrounding region as input.
D. **Quantile-quantile plot of complete topologically associated domains.** Observed versus expected p-values when using all regulatory elements within each HMEC topologically associated domain (Rao et al., 2014) as input.

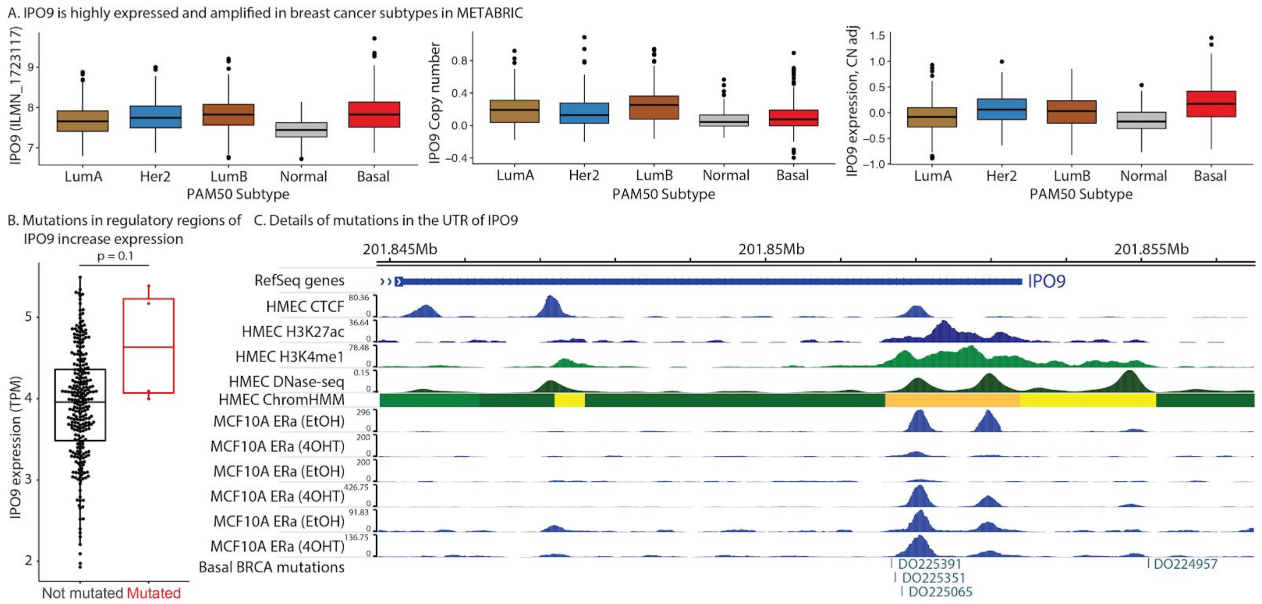


**Supplemental Figure 3: Additional details of IPO9 expression and its locus.**

A. ***IPO9* is highly expressed and amplified in breast cancer subtypes in METABRIC.** Expression, copy number variation, and expression adjusted for copy number variation are presented for each of the PAM50 subtypes in METABRIC.
B. **Mutations in regulatory regions of *IPO9* increase expression.** Increased IPO9 expression in individuals with mutated IPO9 regulatory elements (n=4), relative to non-mutated patients (p = 0.1).
C. **Example of the UTR region harboring additional mutations around IPO9 in breast cancer patients.** Four donors had mutations in this regulatory region, three of which were localized near a shared peak of CTCF, DNase, and histone modification activity.

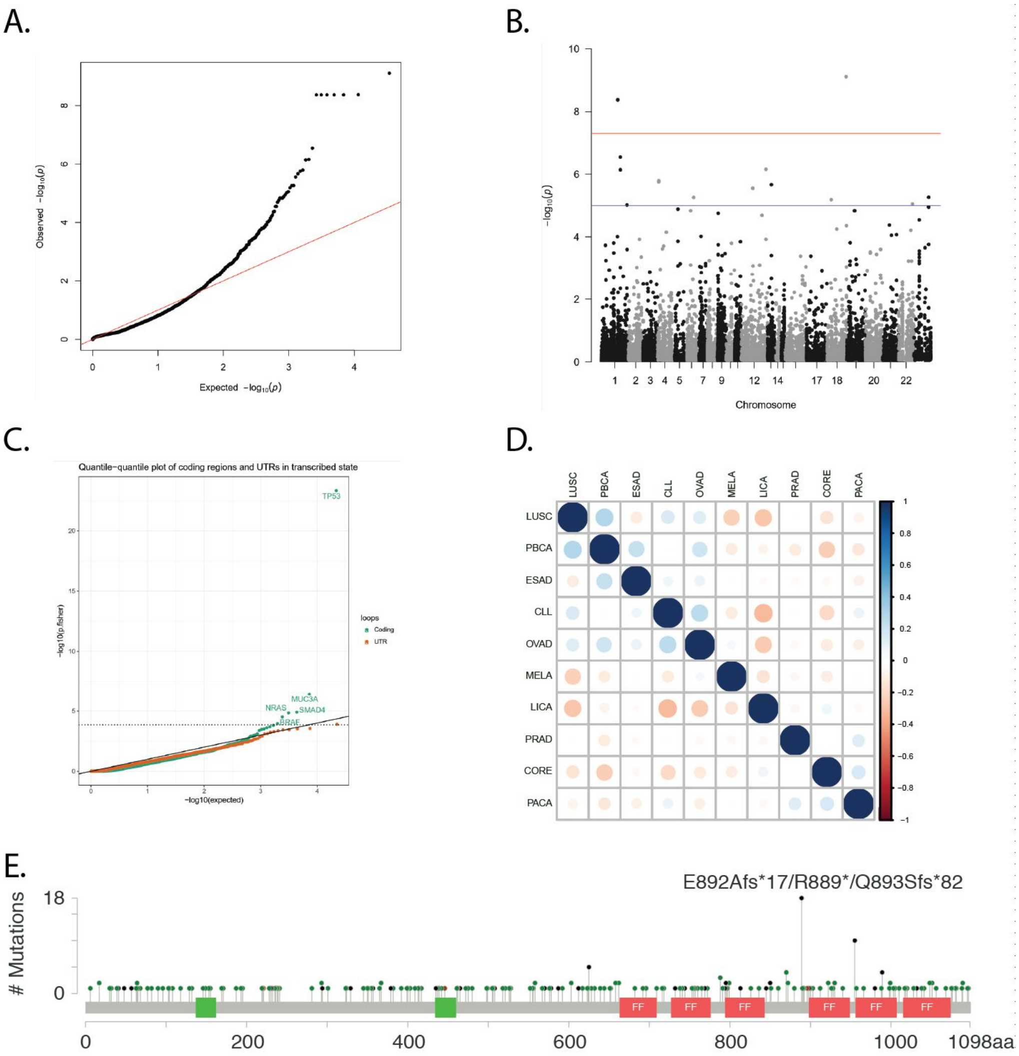


**Supplemental Figure 4: Pan-cancer burden of regulatory element mutations.**

A. **Ovarian cancer enhancer state qq-plot**. Ovarian cancer enhancer chromatin state quantile-quantile plot, illustrating the well calibrated p-values of the analysis.
B. **Ovarian enhancer manhattan plot**. Ovarian cancer enhancer manhattan plot, illustrating the local correlation structure of adjacent genes with respect to their regulatory mutations.
C. **Pan-cancer coding mutation qq-plot.** Coding and UTR mutation qq-plot, showing a substantial increased burden at *TP53* and lack of enrichment over expectation in actively transcribed UTRs.
D. **Correlation of burden across cancer types**. Correlation of mutational burden effects across cancer types, suggesting most burden is not shared across cancer types.
E. **Lollipop plot of coding mutations in *TCERG1***. Lollipop plot of *TCERG1 (Bailey et al., 2018; Gao et al., 2013)* showing two close-by regions of high mutational burden across cases from The Cancer Genome Atlas.

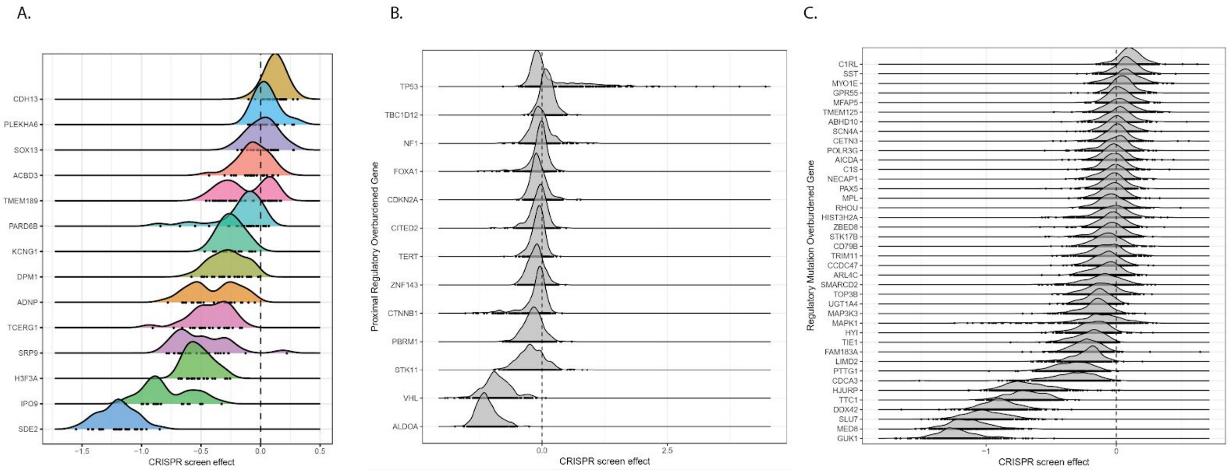


**Supplemental Figure 5: CRISPR growth effects of pan-cancer putative driver genes.**

A. **Growth effect in breast cancer genes among breast cancer lines from Avana.** Detailed growth effect of recurrent breast cancer regulatory genes among breast cancer cell lines from Avana. Each distribution is the effects observed for one of the enriched genes, and dots indicate the individual line effects.
B. **Growth effect of genes with recurrent promoter mutations.** Observed growth effects of genes found to have recurrent promoter mutations in the pan-cancer screen.
C. **Growth effect in Avana for genes with pan-cancer regulatory mutations**. Distributions of effect sizes across cell types for each of the recurrently mutated regulatory genes.

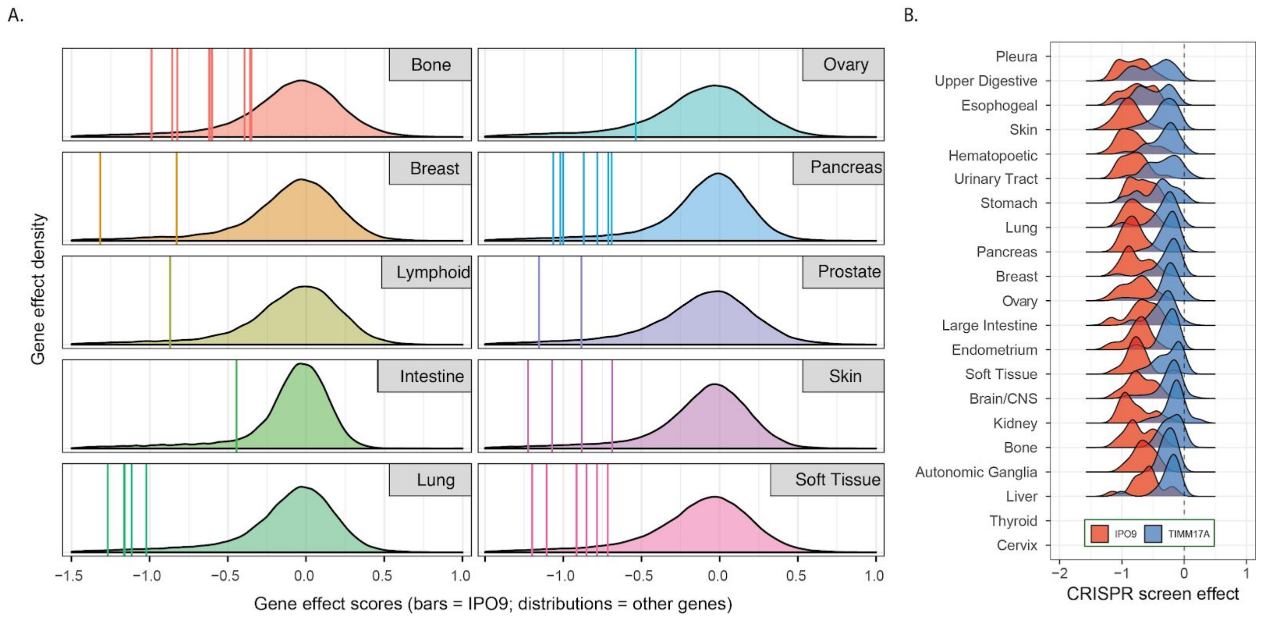


**Supplemental Figure 6: CRISPR growth effects of IPO9 and its locus**

A. **Breakdown of *IPO9* gene effects across cancer types in GeCKOv2**. Independent GECKO screen results from Project Achilles. Vertical lines represent estimates of effect of *IPO9* on cell line growth for each of the 47 cell lines, and the distribution is the observed effects of other genes.
B. **Effects of *TIMM17A* and *IPO9* across tissues.** Comparison of effect of knockout of *IPO9* versus *TIMM17A* (the gene at *IPO9* locus with the second largest growth effect) within cancer type groups in the Avana screening data. Only in cancers isolated from pleura or upper digestive tract is *TIMM17A* similar in gene effect score to *IPO9*.

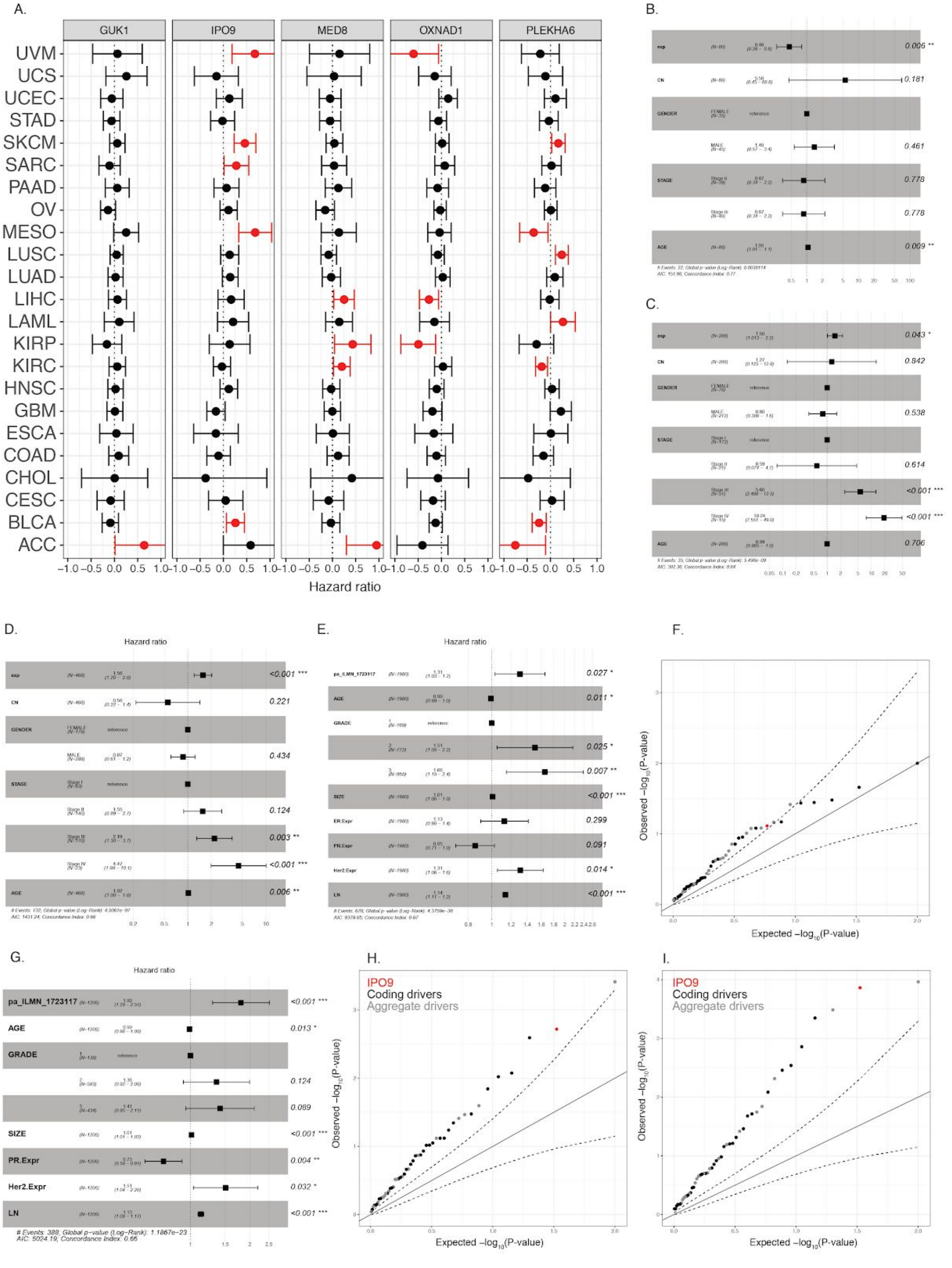


**Supplemental Figure 7: Cox proportional hazard analysis of the association between candidate driver gene expression and survival.**

A. **Hazard forest plot for effect of *GUK1*, *IPO9*, *MED8*, *OXNAD1*, and *PLEKHA6* expression on relapse-free survival in 33 cancer types from TCGA**. Full hazard plot for all five tested genes across the 33 cancer types that comprise the PanCanAtlas.
B. **Hazard forest plot for OXNAD1 among uveal melanoma cases (TCGA)**. Hazard ratio for *OXNAD1* expression adjusted for stage, gender, age, and copy number in the TCGA uveal melanoma dataset.
C. **Hazard forest plot for MED8 among kidney papillary cases (TCGA)**. Hazard ratio for *MED8* expression adjusted for stage, gender, age, and copy number in the TCGA papillary kidney carcinoma dataset.
D. **Hazard forest plot for IPO9 among melanoma cases (TCGA)**. Hazard ratio for *IPO9* expression adjusted for stage, gender, age, and copy number in the TCGA melanoma dataset.
E. **Hazard forest plot for IPO9 expression among all cases (METABRIC)**. Hazard ratio for *IPO9* expression adjusted for stage, grade, age, copy number, and ER, PR and Her2 expression in METABRIC.
F. **QQ-plot for overall survival**. Quantile-quantile plot of breast cancer coding (black) and non-coding (grey) genes in predicting overall survival across all METABRIC cases. IPO9 highlighted in red.
G. **Hazard forest plot for IPO9 in luminal cases (METABRIC)**. Hazard ratio for *IPO9* expression adjusted for stage, grade, age, copy number, and PR and Her2 expression across luminal breast cancers.
H. **QQ-plot for overall survival (luminal)**. Quantile-quantile plot of breast cancer coding (black) and non-coding (grey) genes in predicting overall survival across luminal METABRIC cases. IPO9 highlighted in red.
I. **QQ-plot for cancer-specific survival (luminal)**. Quantile-quantile plot of pan-cancer regulatory genes in predicting cancer specific survival across all METABRIC cases.
